# Between-subject correlation of heart rate variability predicts movie preferences

**DOI:** 10.1101/2020.10.12.335646

**Authors:** Tsz Yan So, Man Yi Erica Li, Hakwan Lau

## Abstract

We introduce a novel and simple method for assessing audiences’ emotional responses to audiovisuals (e.g. films). Viewers (*N*=21) watched movies and TV commercials from different genres while wearing photoplethysmography (PPG) optic sensors on their wrists. Heart rate variability (HRV) synchrony was observed among the audience. Based on this between-subject synchrony measure, we identified emotionally arousing segments from the materials. New participants (*N*=24; *N*=16) were then invited to watch these identified segments along with some randomly selected segments as control; they reported that the former was more engaging (effect size *w*=.67; *w*=.5). This finding was confirmed in an online study with a larger cohort (*N*=300). While some specific effects varied depending on movie genre or gender, HRV-based editing generally performed better than the control. These findings suggest that HRV synchrony can be used as a new tool for audience psychology, and potentially also for automatically creating short trailers out of movies in a principled manner while taking into account the human perspective.

## Introduction

There are increasingly more and more media forms for visual storytelling, be it theatrical films, videos on demand, or TV commercials, for a wide range of goals such as entertainment, education, and marketing. Cognitive film theorists proposed using schémas, inferences, hypotheses, and assumptions in film viewing to structurally and scientifically observe audiences. They concluded that audiences were goal-oriented but non-conscious in the process of making sense of the film narratives [1–5]. In other words, the audience actively comprehends the stories and naturally gains emotional experience through the perception of images, dialogues, and audio effects.

Hollywood commonly uses test screenings to gauge audience responses before the official release of a film [6]. Audience feedback is referenced by the producers to improve the film by re-editing or re-shooting. Nonetheless, self-reporting surveys are subjective and introspective, and lack standardized measurements of audience engagement. It is also difficult to trace the audience’s responses continuously since the feedback is often collected only on a few specific scenes.

Neural measures have arisen as an objective measure of emotions in cognitive psychology. Hasson and colleagues coined the term “*Neurocinematics*”, focusing on bringing cognitive neuroscience and film studies together to understand how the viewer’s brain activity is evoked by audiovisuals [7,8]. Inter-subject correlation (ISC) was used to analyze the similarities in the spatiotemporal response across the audience’s brains, and regions associated with the most primitive stages of sensory processing always responded similarly across individuals watching the same video clip [7]. He then suggested filmmakers use ISC as a quantitative neuroscientific assessment of the engaging power of their productions.

In recent years, there is an increasing interest in the study of HRV and the autonomic correlates of emotions. Heart rate was found to accelerate with certain emotions, such as sadness, anger, and fear, and decelerate with disgust [9]. A quadratic relationship was found between high-frequency (HF) HRV and positive emotions associated with safeness and contentment. Different kinds of positive emotions can be characterized by qualitatively distinct profiles of autonomic activation [10]. This inspires us to look into the association between the ISC of HRV and movie preferences.

We hypothesize that: (1) there are detectable similarities in the audience’s HRV during the viewing of audiovisual materials of different genres, and (2) new audiences would have more engagement toward the edited clips according to previous audience’s emotional arousals indicated by their HRV.

## Study 1: HRV data collection

### Material and methods

In Study 1, we collected HRV responses, survey responses, and facial expression recordings from an audience while they were viewing 4 different genres of movies and commercials. We aimed to detect if any HRV synchrony during the viewing can be observed.

As depicted in the schematic diagram in Fig 1, twenty-one participants (nine males and twelve females) watched a TV commercial compilation and three movie clips in a 105-seat screening room with THX 5.1 Surround Sound System and HD projector. This five-hour-long experiment included four separate viewings of different dramatic elements: Movie A - a Thai commercial compilation (comic), Movie B - the tragic movie *Roma* [11], Movie C - the cerebral sci-fi movie *2001: A Space Odyssey* (abbr. *Space Odyssey*) [12], and Movie D - the action movie *Mission Impossible: Rogue Nation* (abbr. *Mission Impossible*) [13].

**Fig 1.**
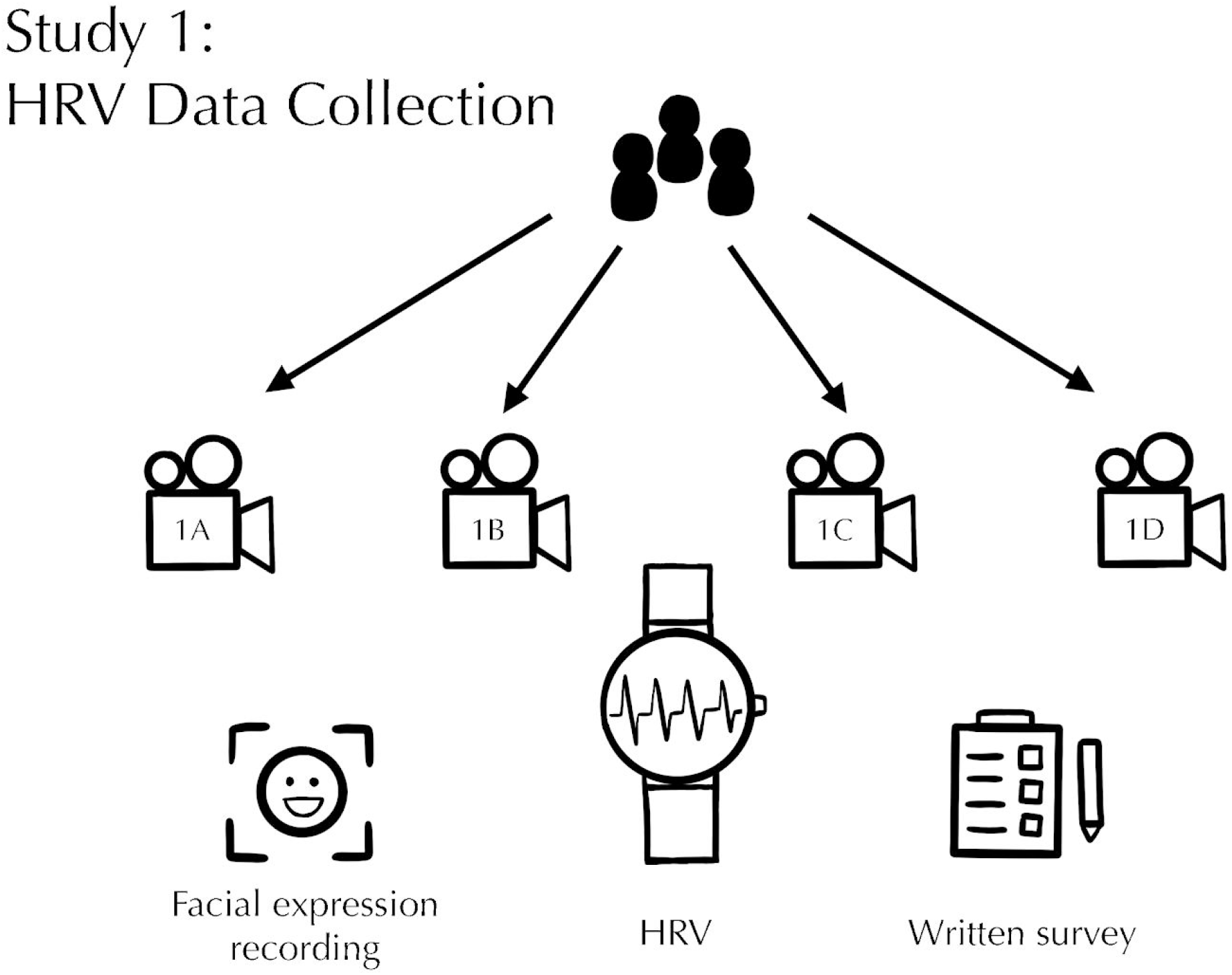
HRV data collection process in Study 1. Schematic diagram of the experimental procedure: participants watched four movies (A, B, C, and D) in sequence while HRV data was collected and facial expressions were recorded.

The participants’ age ranged from 16 to 59 years (*M*=38.09, *SD*=14.02) and were from different backgrounds including businessmen, homemakers, retirees, and students. During the ten min-breaks in between viewings, refreshments and drinks were offered and smoking was allowed, but these activities were strictly prohibited during the viewings.

### HRV data acquisition and processing

Pulse signals of each participant were picked up from their wrists and recorded by Upmood (S1 File), a wearable emotion tracker/mood sensor that collects biodata [14]. The PPG sensor embedded in the device enables a non-intrusive way to measure HRV by analyzing the power of the pulse signal and calculating the different indices such as HF (high frequency), LF (low frequency), LF/HF ratio, and VLF (very low frequency).

To standardize the performance of all Upmood devices, only android mobile phones were used. All phones were either owned by or lent to the participants and were installed with the Upmood app which enables the transmission of data from the PPG sensors to the users’ mobile phones via Bluetooth. All data were transmitted to and collected by Upmood servers in the Philippines. Since our study interest is solely on HRV, only raw data such as timestamps and PPI (P-P intervals, the intervals between the P waves due to atrial depolarization) were used in our analyses.

The Upmood PPG device recorded the participants’ PPI values for approximately four hours, from 14:40 to 18:30 local time. After the experiment, timestamps and raw PPI values in comma-separated values (CSV) format are imported into Python [15] via Pandas [16]. Participants with their first timestamp recorded after 14:40 or their last timestamp recorded before 18:30 were excluded due to data loss. The PPI values were recorded in batches approximately every 90 seconds prior to the recorded timestamp. Since the device was set up to automatically throw out noisy data, linear interpolation with time intervals of 250ms was used to fill in missing data when the PPIs do not add up to the correct timestamps. Using SciPy [17], we applied univariate splines of degree three to fit the data. This is to ensure that the data is locally polynomial to allow derivatives to be taken.

To calculative HRV, we take the absolute values of the derivative from the fitted PPI curves. A moving average of every 100 data points was taken to smooth out the curves. Then, using Scikit-learn [18], a min-max normalization was performed on individual data so that all HRV values were between zero and one for each participant.

### Facial expression data acquisition

Video recording of the participants’ facial expressions was taken at two angles by two SONY HXR-NX30 digital video cameras. All participants were included in the front shot, while five male and five female participants in the first two rows were in the side shot taken at about forty-five degrees to the left of the audience. Two digital clocks were in the shots to indicate the actual time so that the timestamp of the HRV can be synchronized with the audience’s facial reaction and the visual frame of the stimuli. The sound in the screening room was recorded, so that the soundtrack of stimuli can synchronize with corresponding visual frames.

### Questionnaire data acquisition

Participants responded to a written survey (S2 File) upon each viewing. The questionnaire consists of five parts, of which the first part collects personal information including gender, age, and occupation, while the rest are about audiences’ subjective feelings towards the stimuli. For each clip shown, participants had to rate it based on a Likert scale from one to ten and describe the scenes they thought were the happiest, the saddest, the most memorable, and the most boring. The participants also had to indicate if they had watched to clip before.

This research has been approved by the Departmental Research Ethics Committee, Department of Psychology, The University of Hong Kong under the project title “Audience Psychology: Measuring Emotional Response of Audience based on Heart Rate Variability”. Written consent has been obtained from in-person participants and online consent has been obtained from online participants.

### Results

#### HRV synchrony observed in PPI derivatives

After data exclusion due to data loss, we have PPI data from seventeen participants (8 males, 9 females) from 16 to 59 years (*M*=39.23, *SD*=13.69). By taking the derivatives of the raw PPI data, we obtain the normalized HRV graphs for each participant, and those of the first four participants are shown in Fig 2. We can see that there were synchronized spikes, both peaks, and troughs, throughout the five-hour-long study. There was also a synchronized plateau during the forty-five-minute long break and “calm down” period between Movie C and D, at timestamps 17:06-17:51. There was an immediate sharp rise in HRV along with great fluctuation during the transition period from “calm down” time to the viewing of Movie D, *Mission Impossible*, from timestamp 17:51 till the end of the study. By inspection, there was also a difference in the degree of synchrony by genres. Movie A (comic) and Movie D (action) produced stronger HRV synchrony than Movie B (tragic) or Movie C (cerebral sci-fi).

**Fig 2.**
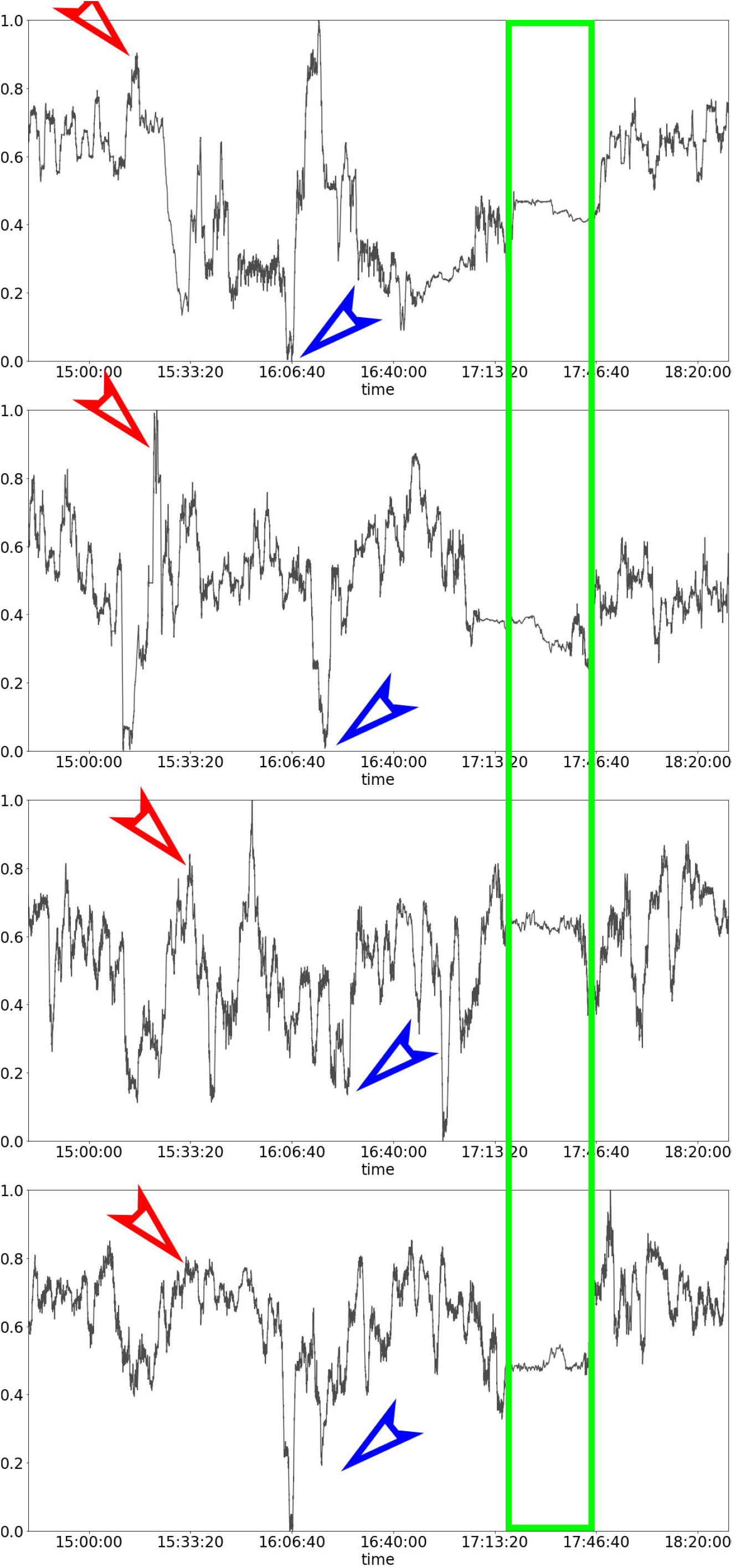
Normalized HRV graphs of four of the participants in the five-hour-long Study 1. The strongest synchrony was marked by the longest plateau (enclosed by the green rectangle) that was produced by the “calm down” period. The graphs also show that synchronized upward spikes (red arrows) and downward spikes (blue arrows) exist.

To quantify this observed synchrony, we computed the Pearson correlation coefficients between pairs of participants (S3 File). 85 out of 136 (63%) correlation coefficients (*N*=55185) were significantly positive, with 73 (54%) of the coefficients highly significant (*p*<.001), 9 (7%) of those moderately significant (*p*<.01), and 3 (2%) of those weakly significant (*p*<.05).

#### HRV corresponds to survey results

The general satisfaction was measured based on a Likert scale from one to ten from the written survey. Movie A (comic) has a mean score of 7.86 (*SD*=1.01). Movie B (tragic) has a mean score of 8.06 (*SD*=1.84). Movie C (cerebral sci-fi) has a mean score of 5.90 (*SD*=2.60). Movie D (action) has a mean score of 7.81 (*SD*=1.12). High arousal points reflected by the mean HRV (Fig 3) correspond to a lot of exciting moments in the clips which were also the participants’ choice of “the happiest, the saddest and the most exciting parts” in the written survey, while low arousal points correspond mostly to calm and sometimes sad moments in the clips. More statistics on the PPI data for each movie can be found in S1 and S2 Figs.

**Fig 3.**
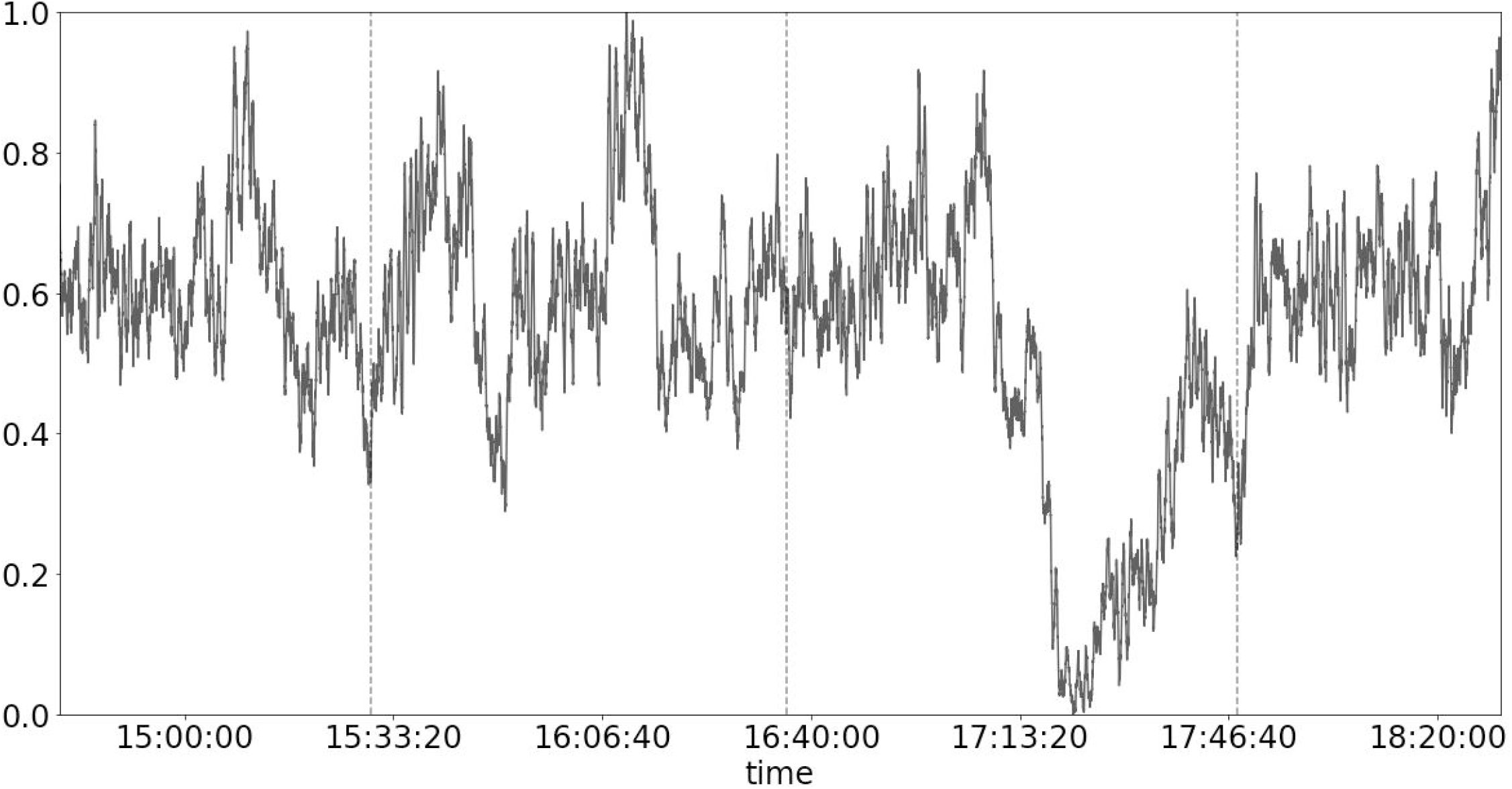
The scale group moving average (over every 100 data points) of heart rate variability over time. Raw pulse-pulse intervals (PPI) values were linearly interpolated and fit with univariate splines of degree three before absolute derivatives were taken. The vertical axis was scaled between zero and one using Min-Max scaling. The vertical dashed lines mark the four sections corresponding to the four movies watched.

## Study 2: Exploratory analysis of audience preferences

### Material and methods

In Study 2, we extracted segments from the audiovisuals based on results from Study 1. We recruited new viewers to watch these edited segments along with randomly selected segments as control over their phones. Chi-square tests were used to determine whether the video clips are equally preferred.

The group averaged HRV (Fig 3) was used to obtain three kinds of video cuts: (1) the ten “most aroused” segments, (2) the ten “least aroused” segments, and (3) ten random segments, from each of the four genres. To ensure that the editing process is double-blinded, the video editor did not know the nature of the cuts and he received only the editing instructions and timestamps from the researcher. Each video cut contains ten segments, and each segment is approximately fifteen seconds long. Hence, the edited clips have a duration of approximately 2.5 minutes each.

Forty participants, who have not participated in Study 1, were given briefings either in person or over the phone. Google links of the comparing video cuts (in MP4 format) were sent to the participants’ mobile phones. To reduce “order effects”, the order of the links was systematically varied for counterbalancing. The participants did not know the nature of the comparison. A group of twenty-four participants (thirteen males and eleven females, from 13 to 60 years, *M*=35.89, *SD*=13.97) was sent the “most aroused” and random clips from one of the four movies. Another group of sixteen participants (ten males and six females, from 22 to 50 years, *M*=34.56, *SD*=8.23) was sent the “most aroused” and “least aroused” clips. Viewers’ engaging preference was asked and replies were sent through mobile phones.

We used an alpha level of .05 for all statistical tests.

### Results

#### Comparison 2A: the “most aroused” vs. random

As summarized in Fig 4, twenty out of twenty-four viewers (83%) said they were more engaged while watching the “most aroused” cut than the random cut. 100% of the viewers chose the “most aroused” cut for Movie B (tragic) and Movie C (cerebral sci-fi), while two-thirds of the viewer preferred the “most aroused” cut for Movie A (comic) and Movie D (action).

**Fig 4.**
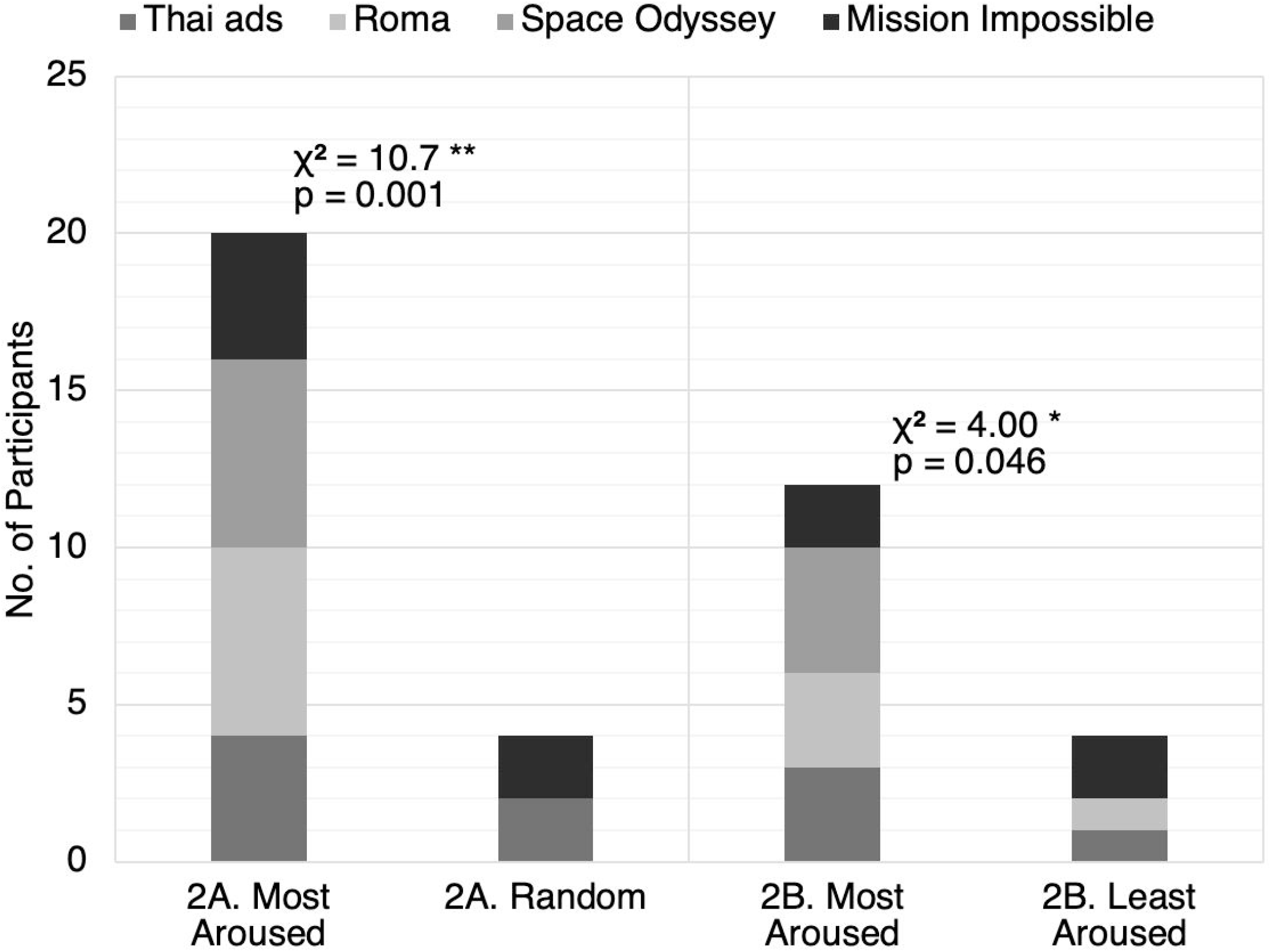
Audience preferences for comparisons 2A and 2B. The bar chart shows the respective number of participants who preferred each video cut. The four movies are represented by different shades. A chi-square goodness of fit test was performed to examine whether participants prefer the “most aroused” cut over (2A) the random cut or (2B) the “least aroused” cut. Both relations were significant. Participants were more likely to prefer the “most aroused” cut over the random cut, χ^2^(1, *N*=24)=10.7, *p*=.001. Participants were also more likely to prefer the “most aroused” cut over the “least aroused” cut, χ^2^(1, *N*=16)=4.00, *p*=.046.

A chi-square goodness of fit test was performed to examine whether the “most aroused” cut and the random cut were equally preferred. The results show that participants were more likely to prefer the “most aroused” cut over the random cut, χ^2^(1, *N*=24)=10.7, *p*=.001 (effect size *w*=.67, power (1-β)=.90). Detailed statistics can be found in <u>S1 Table</u>.

#### Comparison 2B: the “most aroused” vs. the “least aroused”

Twelve out of sixteen viewers (75%) said they were more engaged while watching the “most aroused” cut than the “least aroused” cut. 100% of the viewers chose the “most aroused” cut for movie C (cerebral sci-fi). Three out of four viewers chose the “most aroused” cut for Movie A (comic) and Movie B (tragic), respectively, while the proportions were equally split for movie D (action).

A chi-square goodness of fit test was performed to determine whether the “most aroused” cut and the “least aroused” cut were equally preferred. The preference for the “most aroused” cut is significant, χ^2^(1, *N*=16)=4.00, *p*=.046 (effect size *w*=.5, power (1-β)=.52). Detailed statistics can be found in <u>S2 Table</u>.

This initial exploratory analysis shows promising results and proves the need to recruit more participants and collect more data to gain more statistical power.

## Study 3: Confirmatory Analysis of Audience Preferences

### Preregistration

The confirmatory study has been pre-registered on Open Science Framework (OSF). We hypothesized that “if emotional arousal (derived from previous audience’s HRV) affects movie preferences, then new viewers will be more engaged in the most HRV aroused cut than in the least HRV aroused cut or the random cut from video footage” [19].

### Material and methods

To increase our statistical power, we extended the experiments online. In Study 3, hundreds of participants were recruited to watch these HRV-based edited segments compared to the control. The participants were asked to fill in the DASS-21 [20] and QIDS-SR16 [21] questionnaires. Their strength-of-preference were also collected for meta-analyses.

In addition to the three video cuts from Study 2, we have obtained three more cuts from the group averaged HRV from Study 1: (4) the ten “most synchronous” (defined by the smallest standard deviations among participants) segments, (5) the ten “most aroused” segments based on female viewers only, and (6) the ten “most aroused” segments based on male viewers only.

Each subject will compare two video cuts from one of the four movies, either (A) the “most aroused” and random cuts, (B) the “most aroused” and “least aroused” cuts, (C) the “least aroused” and random cuts, (D) the “most synchronous” and random cuts, or (E) the “most aroused” cuts based on female and based on male.

The experiments were delivered online through Amazon Mechanical Turk (MTurk). We did not impose any demographic restrictions for Comparisons A through D, and fifty participants were recruited for each study. For Comparison E, we requested two groups of fifty participants from MTurk restricted by gender to ensure that we have fifty male and fifty female participants.

After watching the two video clips (order counterbalanced), the participants were asked: “Which video was more engaging” (with four levels of responses - “1st video definitely”, “1st video slightly”, “2nd video slightly”, and “2nd video definitely”). The participants also filled in the DASS-21 [20] and QIDS-SR16 [21] questionnaires (S4 and S5 Files). We grouped the responses “1st video definitely” and “1st video slightly” into a single response of “prefer 1st video”, and likewise for the 2nd video.

We used an alpha level of .05 for all statistical tests.

### Exclusion criteria

The attention of the participants was assessed through the question “What were both videos about”. Participants who failed to answer this question correctly were excluded from our analyses. Any participant who failed to complete the entire study or the questionnaires was also excluded. We continuously rejected participants until fifty eligible responses for each study were received. No participant was allowed to participate in more than one study.

### Demographics

The demographics of the participants of each study are summarized below. Comparison A: 34 males, 14 females, and 2 unstated, from 22 to 59 years, *M*=33.13, *SD*=8.24. Comparison B: 29 males, 19 females, and 2 unstated, from 23 to 60 years, *M*=35.25, *SD*=8.54. Comparison C: 33 males, 15 females, and 2 unstated, from 23 to 66 years, *M*=36.15, *SD*=11.93. Comparison D: 28 males, 18 females, and 4 unstated, from 23 to 74 years, *M*=39.58, *SD*=13.77. Comparison E: 50 males and 50 females, from 18 to 71 years, *M*=38.68, *SD*=11.36.

### Results

#### Comparisons 3A, 3C, and 3D: HRV based vs. random

Participants in studies 3A, 3C, and 3D were asked to rank their preference between the random cut and the “most aroused”, “least aroused”, and “most synchronous” cuts, respectively. As summarized in Fig 5, thirty-one to thirty-two of fifty viewers (62-64%) in each study preferred the HRV based cut more than the random cut.

**Fig 5.**
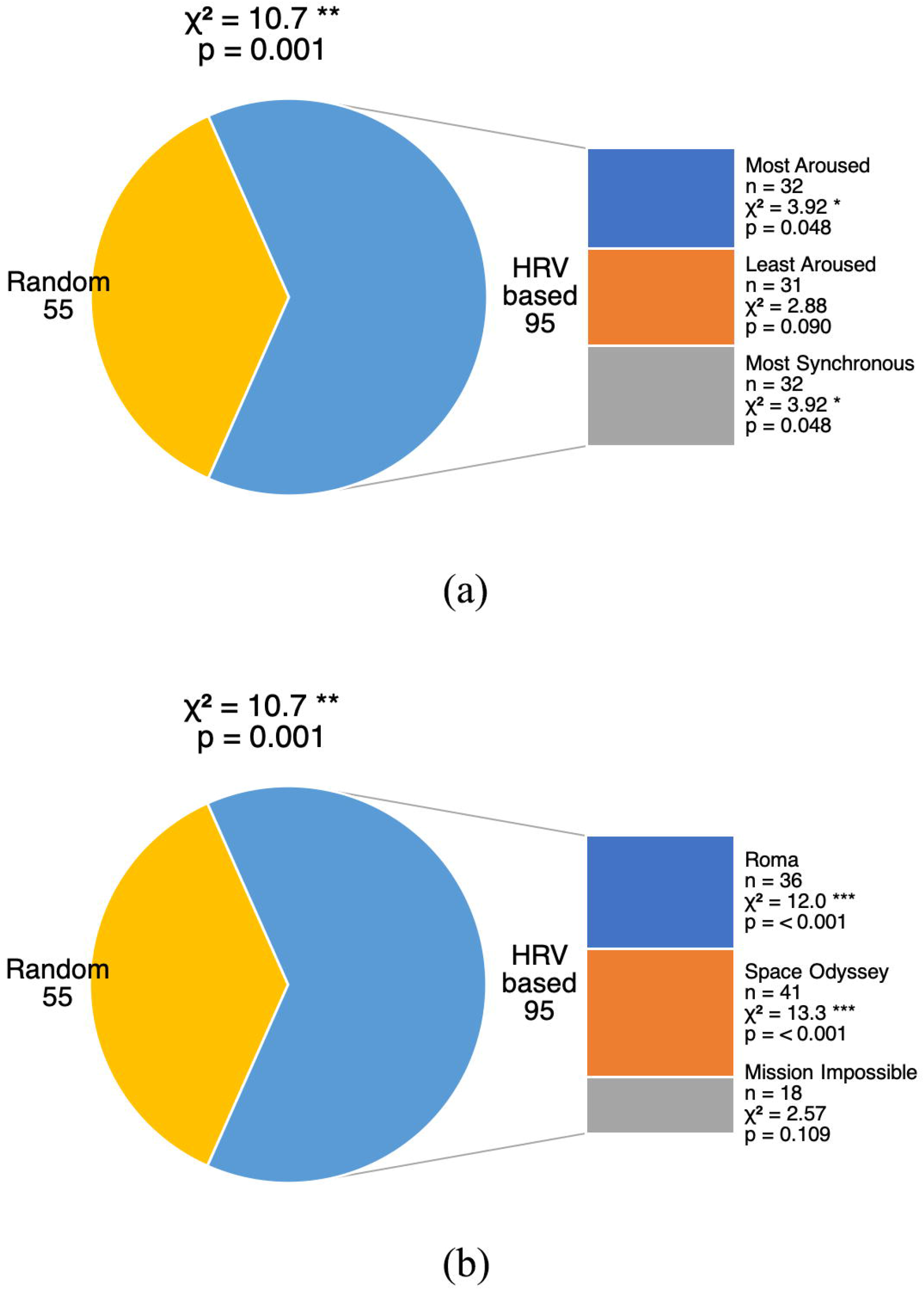
Audience preferences for comparisons 3A, C, and D (a) by study and (b) by movie. The pie chart shows the proportions of participants’ preferences. The sector of HRV based cuts is further broken down to individual bars showing (a) the respective number of participants who preferred each HRV based cut, including (3A) the “most aroused” cut, (3C) the “least aroused” cut, and (3D) the “most synchronous” cut; or (b) the respective number of participants based on the movie watched.

A chi-square goodness of fit test was performed to examine whether the HRV based cuts and the random cut were equally preferred. Overall, participants were more likely to prefer any HRV based cut over the random cut, χ^2^(1, *N*=150)=10.7, *p*=.001 (effect size *w*=.27, power (1-β)=.90).

Looking at the individual studies, only two out of three relations were significant. Participants were more likely to prefer the “most aroused” cut over the random cut, χ^2^(1, *N*=50)=3.92, *p*=.048 (effect size *w*=.28, power (1-β)=.51). Participants were also more likely to prefer the “most synchronous” cut over the random cut, χ^2^(1, *N*=50)=3.92, *p*=.048 (effect size *w*=.28, power (1-β)=.51). The relation was marginally significant between the “least aroused” cut and the random cut, χ^2^(1, *N*=50)=2.88, *p*=.090 (effect size *w*=.24, power (1-β)=.40).

Looking at individual movies, the relation was only significant for two out of three movies. Participants were more likely to prefer the HRV based cuts over the random cut if they watched *Roma* (tragic), χ^2^(1, *N*=48)=12.0, *p*<.001 (effect size *w*=.5, power (1-β)=.93), or *Space Odyssey* (cerebral sci-fi), χ^2^(1, *N*=55)=13.3, *p*<.001 (effect size *w*=.49, power (1-β)=.95). The relation was not significant for *Mission Impossible* (action), χ^2^(1, N=47)=2.57, *p*=.109 (effect size *w*=.23, power (1-β)=.36). Detailed statistics can be found in S3 and S5-S7 Tables.

#### Comparison 3B: the “most aroused” vs. the “least aroused”

The preliminary exploratory data in Study 2B shows a significant preference for the “most aroused” cut over the “least aroused” cut. However, after recruiting more participants for the online study, the effect has diminished.

A chi-square goodness of fit test was performed to examine whether the “most aroused” cut and the “least aroused” cut were equally preferred. The results were not significant overall, χ^2^(1, *N*=50)=2.00, *p*=.157 (effect size *w*=.2, power (1-β)=.29). Moreover, out of our expectation, fifteen out of twenty-one viewers (71.4%) preferred the “least aroused” cut of *Roma* more, and the result was significant regardless of the relatively small sample size, χ^2^(1, *N*=21)=3.86, *p*=.050 (effect size *w*=.43, power (1-β)=.50). Detailed statistics can be found in S4 Table.

#### Comparison 3E: the “most aroused” by female vs. the “most aroused” by male

A chi-square goodness of fit test was performed to examine whether the cut based on female HRV and the cut based on male HRV were equally preferred. As summarized in Fig 6, the results were significant overall, and for two out of the three movies. In general, participants were more likely to prefer the “most aroused” cut based on male HRV, χ^2^(1, *N*=100)=4.84, *p*=.028 (effect size *w*=.22, power (1-β)=.59). Participants were more likely to prefer the cut based on male HRV if they watched *Roma* (tragic), χ^2^(1, *N*=31)=7.26, *p*=.007 (effect size *w*=.48, power (1-β)=.77), or *Mission Impossible* (action), χ^2^(1, *N*=32)=8.00, *p*=.005 (effect size *w*=.5, power (1-β)=.81). The relation was not significant for *Space Odyssey* (cerebral sci-fi), χ^2^(1, *N*=37)=2.19, *p*=.139 (effect size *w*=.24, power (1-β)=.32).

**Fig 6.**
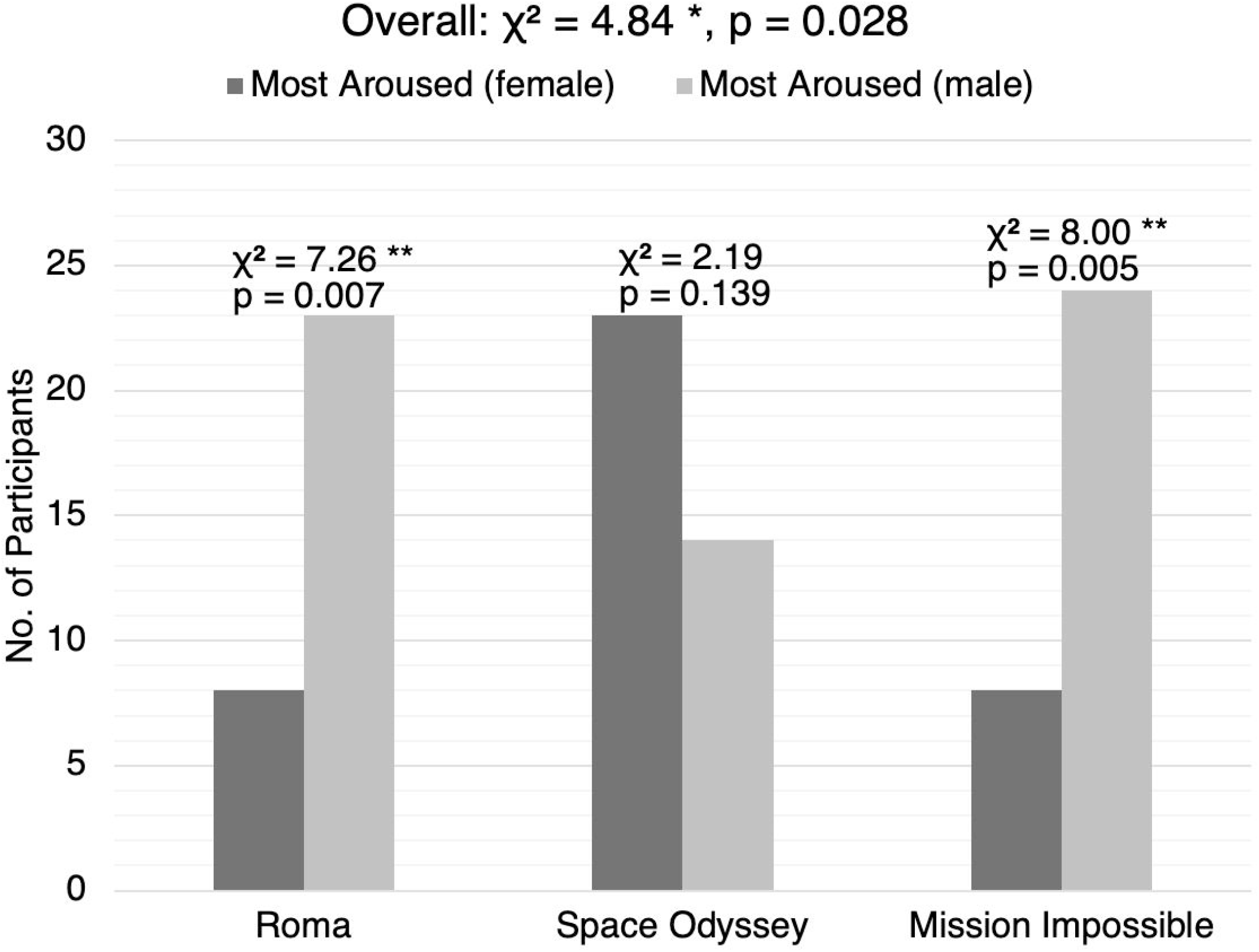
Audience preferences for comparison 3E by movie. The bar chart shows the respective number of participants who preferred the “most aroused” cut based on female HRV and the “most aroused” cut based on male HRV, based on the movie watched.

Detailed statistics can be found in S8 and S9 Tables.

#### Metacognition: Strength-of-preference analyses

In each study, participants were asked to rank their preference for video engagement with four levels of responses: “1st video definitely”, “1st video slightly”, “2nd video slightly”, and “2nd video definitely”. Regardless of the actual movie preference, “definitely” responses were given a strength-of-preference level of one, and “slightly” responses were given a strength-of-preference level of zero.

When grouped by movie, there was a statistically significant difference among groups as determined by one-way ANOVA, *F*(2, 297)=4.59, *p*=.011 (η^2^=.030, power (1-β)=.78). A Tukey post hoc test revealed that the strength-of-preference was significantly higher for *Roma* (tragic; *M*=0.570, *SD*=0.498, *p*=.036, Cohen’s *d*=.34, power (1-β)=.70) and *Mission Impossible* (action; *M*=0.589, *SD*=0.495, *p*=.021, Cohen’s *d*=.38, power (1-β)=.76) compared to *Space Odyssey* (cerebral sci-fi; *M*=0.400, *SD*=0.492). There was no significant difference between the *Roma* and *Mission Impossible* groups (*p*=.963). Detailed statistics can be found in S10 and S11 Tables.

When grouped by comparison condition, there was also a statically significant difference among groups as determined by one-way ANOVA, *F*(4, 295)=4.53, *p*=.001 (η^2^=.057, power (1-β)=.94). A Tukey post hoc test revealed that the strength-of-preference was significantly lower for *“most aroused” vs “least aroused”* (Comparison 3B; *M*=0.440, *SD*=0.501, *p*=.020, Cohen’s *d*=.63, power (1-β)=.88) and *“most aroused” based on female vs male* (Comparison 3E; *M*=0.410, *SD*=0.494, *p*=.001, Cohen’s *d*=.78, power (1-β)=.99) compared to *“least aroused” vs random* (Comparison 3C; *M*=0.740, *SD*=0.443). Details statistics can be found in S12 and S13 Tables.

As for the exploratory analysis, we have not found any significant statistics from the DASS-21 [20] and QIDS-SR16 [21] questionnaires.

## Discussion

The results of Study 1 indicate that some HRV similarities exist among the audience during the viewing of visual storytelling materials, regardless of movie genre. The findings are consistent with our hypothesis. For low-arousal audiovisual materials that were intended to induce calmness between the actual movies, synchronized “plateaus” in the HRV graphs (Fig 2) indicate unanimous steadiness in heart rate responses. For more dramatic audiovisual materials, HRV synchrony is shown in forms of synchronized upward and downward spikes in the HRV graphs. The most observable synchrony in the HRV responses is during the third calm down period, between Movie C and D.

Corresponding to the expected emotional arousal of each movie, we also found HRV dependence on genres. The average HRV is significantly lower (S1 Fig d, Movie A: *t*(1198)=12.9, *p*<.001; Movie B: *t*(1198)=12.3, *p*<.001; Movie D: *t*(1198)=21.4, *p*<.001) for Movie C *Space Odyssey* (dialogue-free cerebral sci-fi), suggesting that boredom, as mentioned in the survey responses of some participants, be another affective state of low arousal-positive valence.

Highly engaging content has the potency of “mind control” through inducing viewers’ neural responses [7,22]. Barnett and Cerf [22] used electroencephalogram (EEG) to measure cross-brain-correlation (CBC) across viewers. Investigating further into viewers’ behavior, they found that ISC correlates positively to the recall of the trailers, and movies with better recall of their trailers have higher average weekly ticket sales [23]. The findings in Study 2 may serve as another piece of evidence of their findings on viewers’ movie preferences. In general, new viewers prefer the HRV based cuts compiled from segments that are either the most or least arousing, or the most synchronous to a previous audience. In other words, the HRV based content is generally more engaging than just random content from a movie, consistent with our second hypothesis.

It is worth noting that our HRV based cuts were entirely based on the “previous audience” but tested on “new audiences”. Thus, the relationship between arousal as reflected in HRV and preferences of content could be established, shedding light on the prediction of future audience’s responses.

Study 1 has reconfirmed the validity of HRV in monitoring emotional arousals, especially during relax and calm affective states, which are of low arousal-positive valence. This explains why HRV tracking by PPG wearables has been widely adopted to monitor heart rate and blood pressure during yoga practices and meditations [24,25].

We also observed that the “most aroused” segments of *Roma* (tragic) had HRV values significantly higher (S1 Fig a, Movie A: *t*(1198)=-38.7, *p*<.001; Movie C: *t*(1198)=-22.4, *p*<.001; Movie D: *t*(1198)=-20.3, *p*<.001) while its “least aroused” and “most synchronous” segments had HRV values significantly lower than the other movies (S1 Fig b, Movie A: *t*(1198)=83.8, *p*<.001; Movie C: *t*(1198)=74.5, *p*<.001; Movie D: *t*(1198)=59.6, *p*<.001; S1 Fig c, Movie A: *t*(1198)=61.6, *p*<.001; Movie C: *t*(1198)=50.4, *p*<.001; Movie D: *t*(1198)=39.9, *p*<.001). The overall range of HRV values of *Roma* was also wider than that of the other movies (S1 Fig). As low HRV could indicate the person is under stress [26] and high HRV could indicate sadness [9], our results suggest that *Roma* could be the more emotionally inducing movie out of the four. The audience seemed to have more extreme emotions when watching *Roma* compared to the other movies.

Another possible explanation for the wider range of HRV could be behavioral expressivity differences in gender or culture. Positive emotion induced behavior such as laughing is generally more socially acceptable than negative emotion induced behavior such as weeping and crying in public. In collectivistic cultures, people may be less willing to express negative emotions than people in individualistic cultures [27], this may apply to our sample of Hong Kong viewers who may not choose to display behavior associated with negative feelings in a shared screening room. Restraining physical actions such as frowning or crying may have an effect upon HRV which reflects physiological reactions.

Out of the three movies we experimented with in Study 3, *Space Odyssey* (cerebral sci-fi) had significantly lower strength-of-preference (S10 and S11 Tables) compared to the other two movies, *Roma* (tragic, *p*=.036), and *Mission Impossible* (action, *p*=.021). While we can look at it as a genre-specific difference, it might be more related to the nature of the film instead. As *Space Odyssey* is generally considered boring to the modern audience [28], it is understandable that the marginal differences between different cuts are small and that the viewers are just not very engaged overall.

While our small sample in Study 2 showed a preference for the “most aroused” cut over the “least aroused” cut, we did not observe the same effect in Study 3 with a larger online sample. These inconclusive results show that the “most aroused” and “least aroused” cuts may be capturing segments from the films that are engaging in different ways. This further confirms that low HRV could be due to stress [26], and stress while watching a film could indicate that the scenes are intense. In particular, by inspection of the segments from *Mission Impossible*, the “most aroused” segments tend to capture the actual fight scenes, while the “least aroused” segments tend to capture the suspense before weapons are drawn.

While we have not observed any statistically significant difference in preference based on the new viewer’s gender, we have observed a genre-specific gender difference in terms of whether we edited the segments based on HRV data from the female or male audience only (Fig 6). The “most aroused” segments extracted from only male HRV data were significantly preferred over those extracted from only female data, for both *Roma* (*p*=.007) and *Mission Impossible* (*p*=.005). For *Space Odyssey*, we observe a non-significant preference (*p*=.139) in the opposite direction, which again reconfirmed that it is just not a very engaging film overall. Coincidentally, the average HRV of the “most aroused” segments from males was also significantly higher (S2 Fig d, *t*(1198)=-15.6, *p*<.001) than that from females for *Mission Impossible*, this could suggest that higher arousal while viewing can predict the viewing preferences of future audiences. While this could suggest that HRV data from males might be more “useful” in predicting audience engagement than that from females, we must be aware that there have been studies showing that heart rate variability is lower in women than men in a healthy population [29]. This gender difference in HRV and its effect on audience engagement should be studied more carefully in the future, possibly by preprocessing data from the two genders separately and performing normalization differently.

Audience psychology requires researching with large samples. However, functional magnetic resonance imaging (fMRI) is expensive and not portable, while EEG obstructs participants’ normal activities and is not easy to administer, meaning they can only be used to test on relatively smaller samples. Although HRV is more commonly derived from R-R intervals (the intervals between the R waves due to ventricular depolarization) measured by electrocardiography (ECG) in the literature [30], HRV derived from PPIs measured by PPG are insignificantly different as shown in experiment [31]. HRV serves as an accessible and scientific tool since PPG devices are easily accessible. It is proposed that future productions and marketing of audiovisual materials can use audience HRV monitoring during test screenings, providing detailed arousal and emotional response records of a large audience.

The utilization of HRV predictions of emotions should not be limited to only movies and TV. Emotional experience from the consumption of any products and services, such as driving a new car, playing a new video game, or even touring a new exhibit in museums can also be potentially studied using HRV in the future.

## Conclusion

HRV can serve as a scientific tool to objectively evaluate the emotion of an audience in the development of audience psychology. More and more recent studies have demonstrated that HRV is an accurate indicator of emotions for certain valence and arousal levels. With wearable fitness devices becoming increasingly popular among consumers and artificial intelligence (AI) getting cheaper and more efficient, HRV measurements are now easily accessible and can create opportunities for filmmakers to systematically edit and improve their productions by technologies in machine learning. The technique we have developed can easily be applied to assess user engagement in similar fields, such as video gaming and music productions. Further studies on the relationship between HRV and different kinds of emotions can also expand its applications outside the creative industries. Perhaps, in the future, HRV monitoring might be able to assist less verbal or facially expressive people, such as young children, patients with autism spectrum disorder (ASD), or even patients with locked-in syndrome, to efficiently express their own emotions and understand others’ emotions.

## Acknowledgments

We would like to express our gratitude for Lo Kit Yi (Charles) of Upmood for providing smartphones, all PPG devices and collecting HRV data for this study, Emperor Movie Production & Emperor Entertainment Group for providing a spacious and comfortable screening room for the participants, Carman Tsang and Helen Wong for helping with the video recording, and Jinpo Yip, Mae Lui and Ken Hui for assisting with the video editing in a professional studio.

## Supporting Information

*Note. * p<*.*05, ** p<*.*01, *** p<*.*001, **** p<*.*0001*

**S1 Fig. Boxplots of the mean absolute derivatives from PPI diagrams of each movie grouped by video cuts**: (a) the “most aroused” cut; (b) the “least aroused” cut; (c) the “most synchronous” cut; (d) the random cut; (e) the female “most aroused” cut; (f) the male “most aroused” cut.

**S2 Fig. Boxplots of the mean absolute derivatives from PPI diagrams of each cut grouped by movies**: (a) Thai commercials; (b) *Roma*; (c) *2001: A Space Odyssey*; (d) *Mission Impossible: Rogue Nation*.

**S1 Table.** Chi-Square Goodness of Fit Test for Comparison 2A (proportions in parentheses).

|  | Thai<br>commercials | Roma | 2001: A Space<br>Odyssey | Mission<br>Impossible:<br>Rogue Nation | Total |
| --- | --- | --- | --- | --- | --- |
| most aroused | 4 | 6 | 6 | 4 | 20 (0.833) |
| random | 2 | 0 | 0 | 2 | 4 (0.167) |
| $\chi^2$ | - | - | - | - | 10.7 ** |
| p-value | - | - | - | - | 0.001 |

**S2 Table.** Chi-Square Goodness of Fit Test for Comparison 2B.

|  | Thai<br>commercials | Roma | 2001: A Space<br>Odyssey | Mission<br>Impossible:<br>Rogue Nation | Total |
| --- | --- | --- | --- | --- | --- |
| **most aroused** | 3 | 3 | 4 | 2 | 12 (0.750) |
| **least aroused** | 1 | 1 | 0 | 2 | 4 (0.250) |
| $\chi^2$ | - | - | - | - | 4.00 * |
| **p-value** | - | - | - | - | 0.046 |

**S3 Table.** Chi-Square Goodness of Fit Test for Comparison 3A.

|  | Roma | 2001: A Space<br>Odyssey | Mission<br>Impossible: Rogue<br>Nation | Total |
| --- | --- | --- | --- | --- |
| **most aroused** | 12 (0.750) | 12 (0.750) | 8 (0.444) | 32 (0.640) |
| **random** | 4 (0.250) | 4 (0.250) | 10 (0.556) | 18 (0.360) |
| $\chi^2$ | 4.00 * | 4.00 * | 0.222 | 3.92 * |
| **p-value** | 0.046 | 0.046 | 0.637 | 0.048 |

**S4 Table.** Chi-Square Goodness of Fit Test for Comparison 3B.

|  | Roma | 2001: A Space<br>Odyssey | Mission<br>Impossible: Rogue<br>Nation | Total |
| --- | --- | --- | --- | --- |
| **most aroused** | 6 (0.286) | 8 (0.444) | 6 (0.545) | 20 (0.400) |
| **least aroused** | 15 (0.714) | 10 (0.556) | 5 (0.455) | 30 (0.600) |
| $\chi^2$ | 3.86 * | 0.222 | 0.0909 | 2.00 |
| **p-value** | 0.050 | 0.637 | 0.763 | 0.157 |

**S5 Table.**
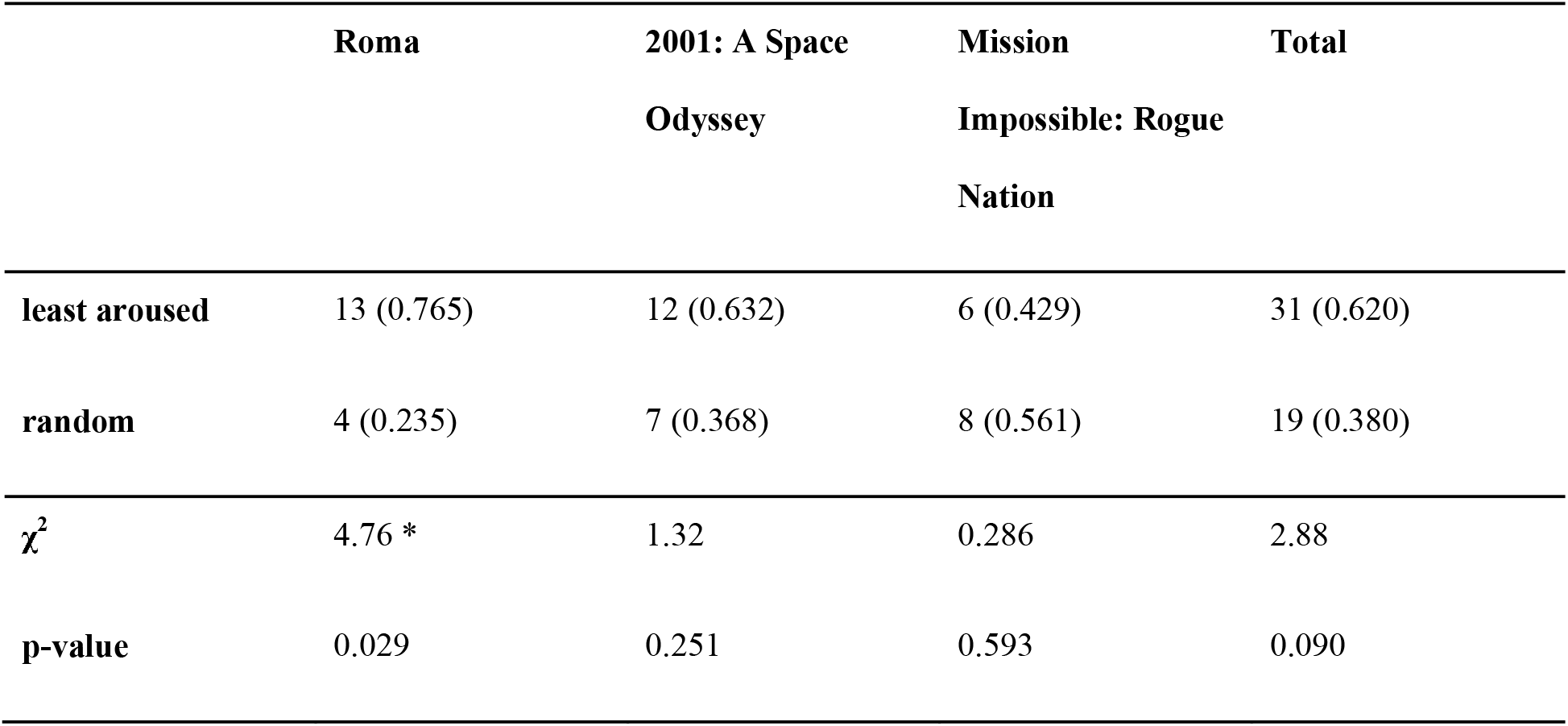
Chi-Square Goodness of Fit Test for Comparison 3C.

|  | Roma | 2001: A Space<br>Odyssey | Mission<br>Impossible: Rogue<br>Nation | Total |
| --- | --- | --- | --- | --- |
| **least aroused** | 13 (0.765) | 12 (0.632) | 6 (0.429) | 31 (0.620) |
| **random** | 4 (0.235) | 7 (0.368) | 8 (0.561) | 19 (0.380) |
| $\chi^2$ | 4.76 * | 1.32 | 0.286 | 2.88 |
| **p-value** | 0.029 | 0.251 | 0.593 | 0.090 |

**S6 Table.** Chi-Square Goodness of Fit Test for Comparison 3D.

|  | Roma | 2001: A Space<br>Odyssey | Mission<br>Impossible: Rogue<br>Nation | Total |
| --- | --- | --- | --- | --- |
| **most synchronous** | 11 (0.733) | 17 (0.850) | 4 (0.267) | 32 (0.640) |
| **random** | 4 (0.267) | 3 (0.150) | 11 (0.733) | 18 (0.360) |
| $\chi^2$ | 3.27 | 9.80 ** | 3.27 | 3.92 * |
| **p-value** | 0.071 | 0.002 | 0.071 | 0.048 |

**S7 Table.** Chi-Square Goodness of Fit Test for Comparisons 3A, 3C, and 3D.

|  | Roma | 2001: A Space<br>Odyssey | Mission<br>Impossible: Rogue<br>Nation | Total |
| --- | --- | --- | --- | --- |
| **HRV based** | 36 (0.750) | 41 (0.745) | 18 (0.383) | 95 (0.633) |
| **random** | 12 (0.250) | 14 (0.255) | 29 (0.617) | 55 (0.367) |
| $\chi^2$ | 12.0 *** | 13.3 *** | 2.57 | 10.7 ** |
| **p-value** | < .001 | < .001 | 0.109 | 0.001 |

**S8 Table.** Chi-Square Goodness of Fit Test for Comparison 3E by movie.

|  | Roma | 2001: A Space<br>Odyssey | Mission<br>Impossible: Rogue<br>Nation | Total |
| --- | --- | --- | --- | --- |
| **female most**<br>**aroused** | 8 (0.258) | 23 (0.622) | 8 (0.250) | 39 (0.390) |
| **male most aroused** | 23 (0.742) | 14 (0.378) | 24 (0.750) | 61 (0.610) |
| $\chi^2$ | 7.26 ** | 2.19 | 8.00 ** | 4.84 * |
| **p-value** | 0.007 | 0.139 | 0.005 | 0.028 |

**S9 Table.** Chi-Square Goodness of Fit Test for Comparison 3E by gender.

|  | female participants | male participants | Total |
| --- | --- | --- | --- |
| **female most**<br>**aroused** | 19 (0.380) | 20 (0.400) | 39 (0.390) |
| **male most aroused** | 31 (0.620) | 30 (0.600) | 61 (0.610) |
| $\chi^2$ | 2.88 | 2.00 | 4.84 * |
| **p-value** | 0.090 | 0.157 | 0.028 |

**S10 Table.** Strength-of-preference grouped by movie.

|  | movie | N | Mean | SD |
| --- | --- | --- | --- | --- |
| strength-of-preference | Roma | 100 | 0.570 | 0.498 |
|  | 2001: A Space Odyssey | 110 | 0.400 | 0.492 |
|  | Mission Impossible: Rogue Nation | 90 | 0.589 | 0.495 |

*One-Way ANOVA: $F=4.59$ \*, $df1=2$ , $df2=297$ , $p=.011$*

**S11 Table.** Tukey Post-Hoc Tests on Strength-of-preference grouped by movie.

|  |  | Roma | 2001: A Space<br>Odyssey | Mission Impossible:<br>Rogue Nation |
| --- | --- | --- | --- | --- |
| **Roma** | Mean difference | - | 0.170 * | -0.0189 |
|  | p-value |  | 0.036 | 0.963 |
| **2001: A Space**<br>**Odyssey** | Mean difference | - | - | -0.1889 * |
|  | p-value |  |  | 0.021 |
| **Mission Impossible:**<br>**Rogue Nation** | Mean difference | - | - | - |
|  | p-value |  |  |  |

**S12 Table.** Strength-of-preference grouped by comparison condition.

|  | comparison | N | Mean | SD |
| --- | --- | --- | --- | --- |
| **strength-of-preference** | most aroused vs. least aroused | 50 | 0.440 | 0.501 |
|  | most aroused vs. random | 50 | 0.600 | 0.495 |
|  | least aroused vs. random | 50 | 0.740 | 0.443 |
|  | most synchronous vs. random | 50 | 0.480 | 0.505 |
|  | female most aroused vs. male most aroused | 100 | 0.410 | 0.494 |

*One-Way ANOVA: $F=4.53^{**}$ , $df1=4$ , $df2=295$ , $p=.001$*

**S13 Table.**
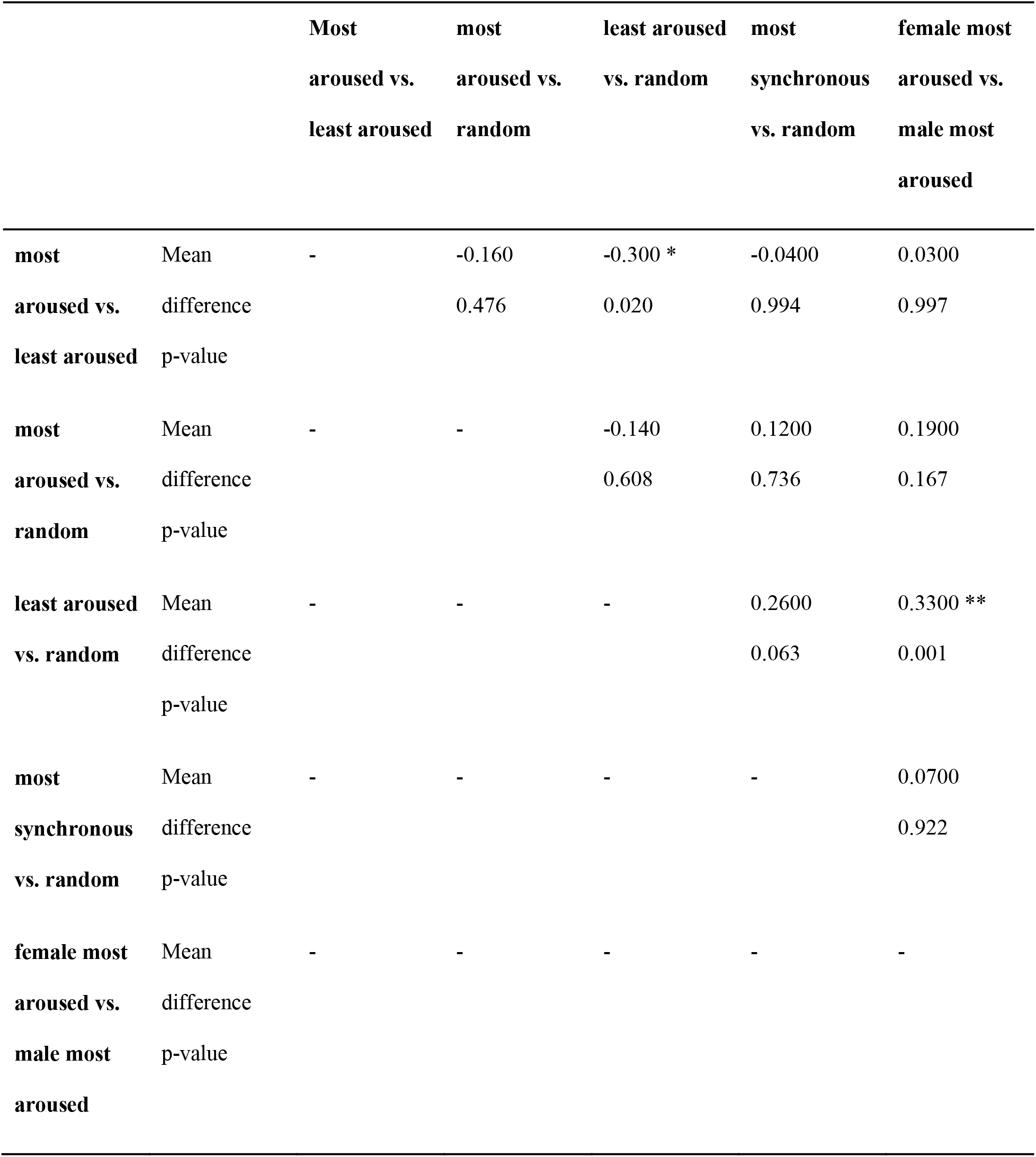
Tukey Post-Hoc Tests on Strength-of-preference grouped by comparison.

**S1 File. Upmood User Guide**.

**S2 File. Study 1 Questionnaire**.

**S3 File. Correlation matrix of HRV across participants**.

**S4 File. DASS-21 Questionnaire**.

**S5 File. QIDS-SR16 Questionnaire**.

## Notes

https://osf.io/xj8gt/

### Competing Interest Statement

The authors have declared no competing interest.

