## Supplementary figures and images for "Between-subject correlation of heart rate variability predicts movie preferences"

### S1 Fig

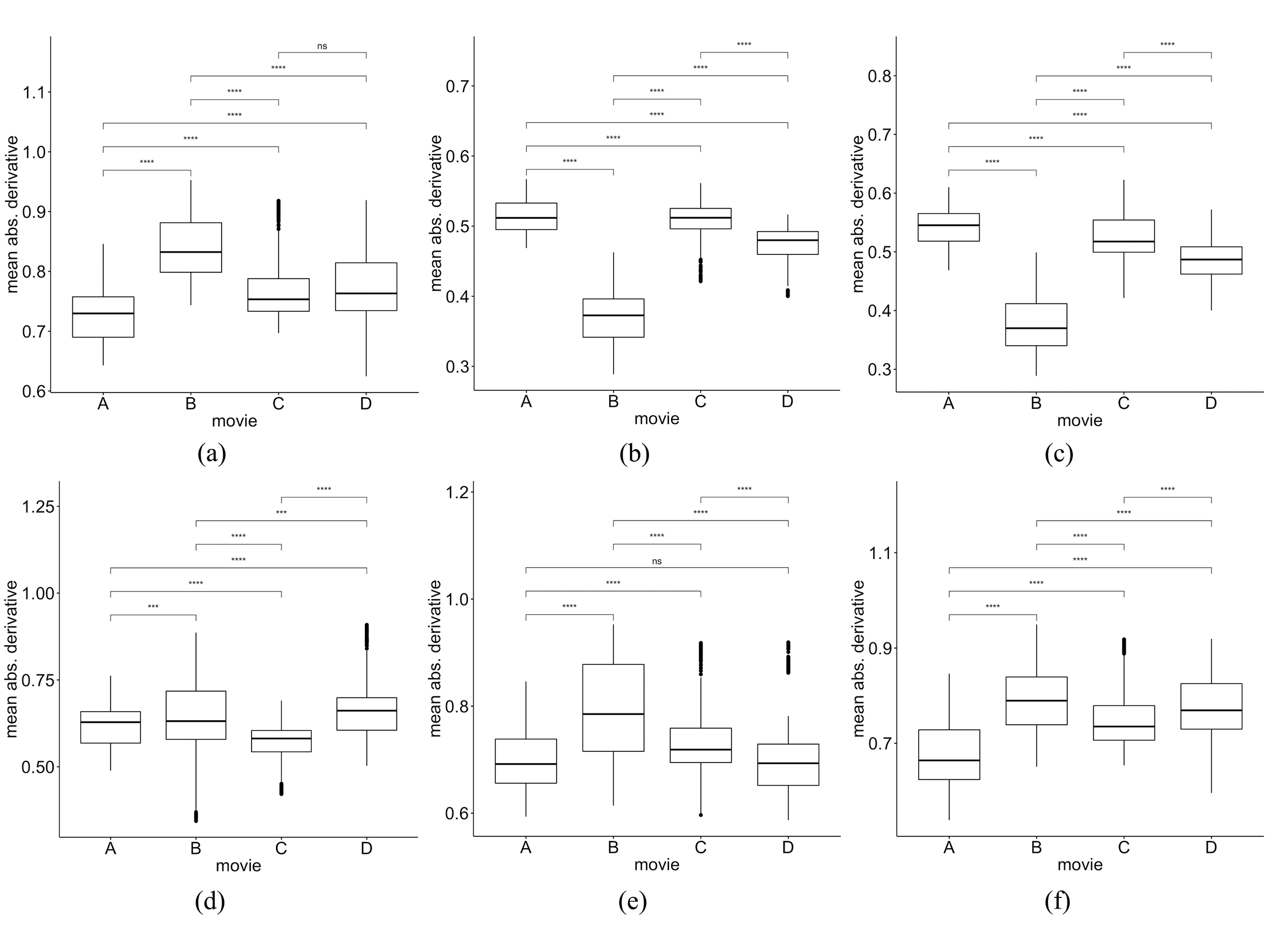

### S2 Fig

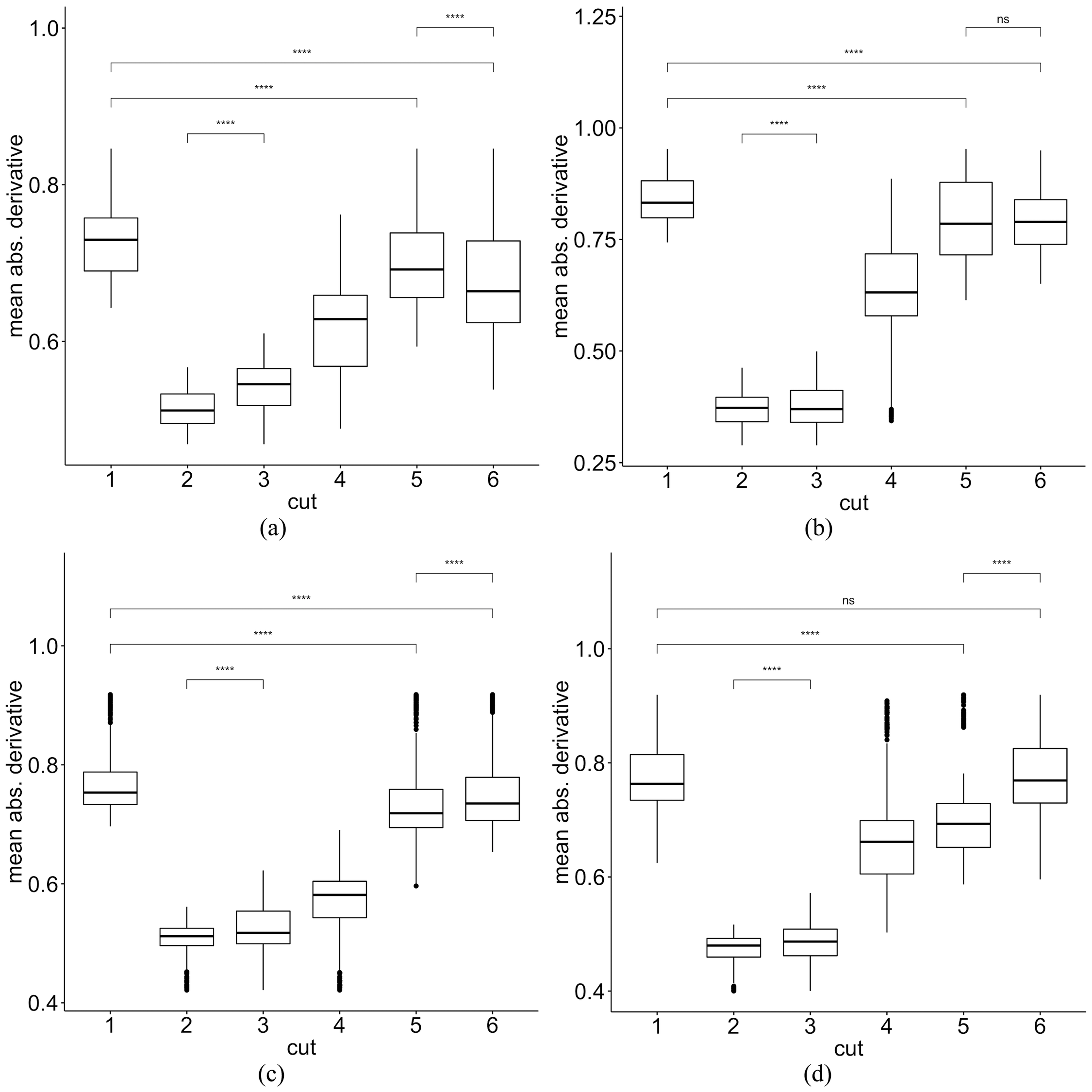
