## Supplementary material for "Between-subject correlation of heart rate variability predicts movie preferences": S1 File

#### Charging Upmood Band

1. Plug the magnetic charger into a USB outlet.
2. Attach the magnetic charger onto the back side of the Upmood device from the right.
3. Once its attached, device will like up in RED. Light turns BLUE when when its fully charged.

Note: Do not force the magnetic charger onto the conductive points. Charger would attach magnetically with the correct orientation.

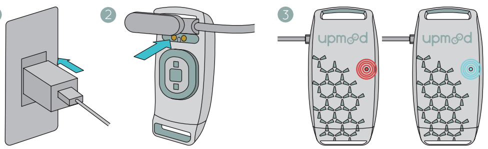

#### Turning the Band ON/OFF

1. Touch and hold onto the logo of the Upmood device for 3 secs to turn ON device.
2. Touch and hold onto the logo of the Upmood device for 3 secs to turn OFF device.

Note: Once the band is connected to the device the ON/OFF is disabled. To able it again, disconnect the band from the app.

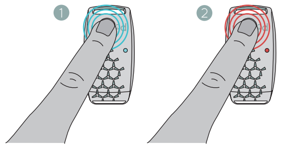

#### Turning the Band ON/OFF

1. Place the band on your wrist
2. Loop and hook the band from the inside through the hole found at the top of the band head.
3. Pull the strap and adjust it to fit comfortably on your wrist.
4. Secure the band by pressing the Velcro strips together.

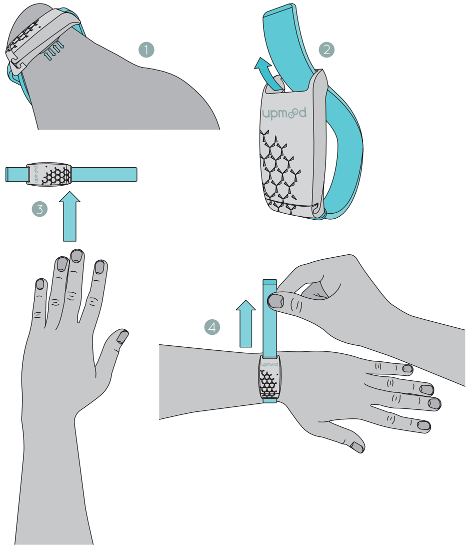

#### Connecting Upmood Band to App

1. Tab on to device on the Upmood App.
2. Scan for Upmood band.
3. Tab to connect.
4. Press disconnect to disconnect band.

Note: If disconnect is not responding please go to Account > Settings and deactivate Autoreconnect band.
The Upmood app is free and available on Google Play and App Store!

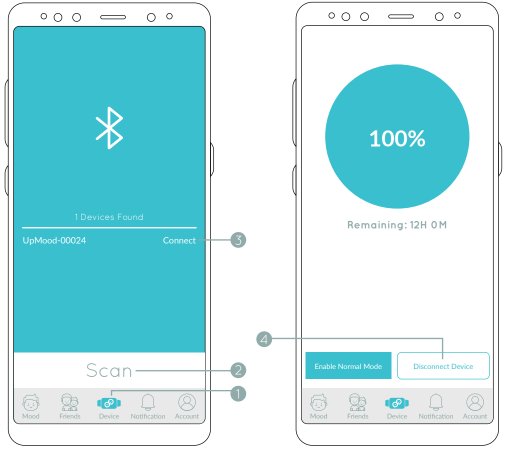


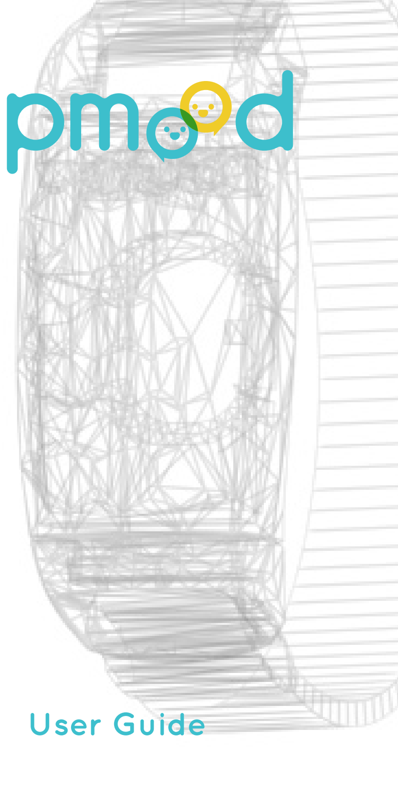

### upmood

#### User Guide

#### Package Inclusion

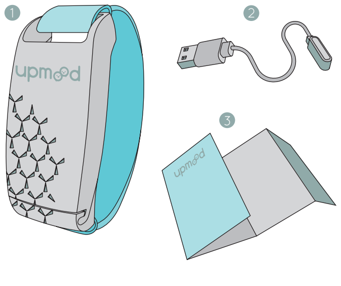


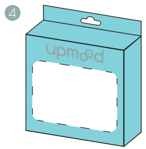


1. 1x Upmood band
2. 1x USB
3. 1x user manual
4. 1x box

#### Specifications

| | |
|---|---|
| **General Specs** | Compatible with all iOS and Android devices<br>PPG Sensor |
| **Strap** | Interchangeable polyester strap<br>Adjustable size: 1” to 9” (diameter) |
| **Battery** | Li-polymer battery<br>48 hrs standby time<br>12 hrs always on time<br>1 hour charging time |
| **Connectivity** | Bluetooth |
| **Waterproof** | IP68 |

**Warning:**
(1) Upmood is not a medical device. It’s simply a tracker to detect your emotions. (2) If you are experiencing any allergic reaction or skin irritation from wearing the band, stop using it and consult a doctor. (3) Do not use the band while it is charging. (4) Do not disassemble, or attempt to repair the band yourself. (5) Do not wash Upmood with soap or any chemical. To clean the strap, remove it from the Upmood device.

#### Parts of the Band

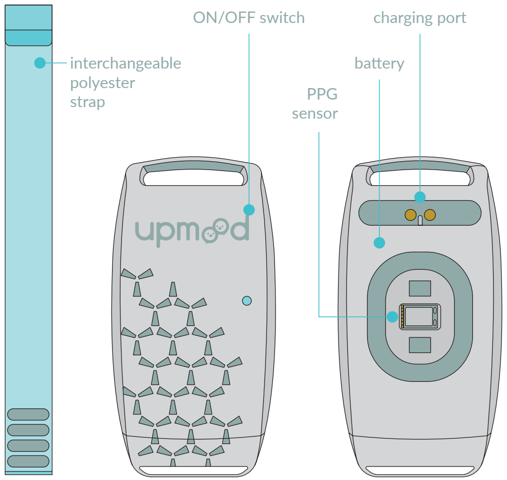


**Warranty:** Upmood provides a one (1) year warranty for its products, valid from the date of purchase. Warranty is rendered invalid and void upon the following:
(1) Damage on the battery or external parts subject to normal customer use. (2) Damage resulting from misuse, alteration, tampering or non-compliance with the precautions outlined in Upmood’s user manual. (3) Damage, losses or defects directly or indirectly related to the product. (4) Products purchased, repaired, or replaced from/by any unathorized third party, unless otherwise stipulated by local law. (5) Products with its serial number been altered, removed or tampered.

The warranty will only be provided for products purchased from authorized retailers.

**Troubleshooting:** When you are experiencing technical difficulties with your Upmood product, try these troubleshooting techniques:
(1) If you are unable to pair Upmood with your device, check if your bluetooth connection is enabled. (2) If the band is unable to detect your emotion, check if any body hair and/or tattoo is blocking the sensor. (3) If the band still doesn’t work, try wearing it on your other wrist.

#### Emoji Explanation

Upmood uses the parameter of emotional arousal, duration and stress level to result in 11 different states. These states are then labeled with text that can be best described the mood but they do not represent the a scientific/diagnostic definition.

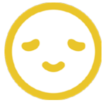

**Calm** is a mental state of peace, tranquility, and relaxation. An Upmood user who has low to moderate stress levels and a steady emotional state throughout the day will be considered in a calm mood.

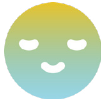

**Zen** is a holistic approach on total state of focus that is achieved when an Upmood user’s mind and body has achieved a meditative state for a period of time, amidst the presence of stress and emotional arousal.

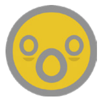

Upmood will most likely display the **Tense** emotion when situations such as contemplation about taking risks, control of a situation is lost, or fear of things going wrong, arises.

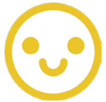

**Pleasant** is an emotion evoked by positive moments. It is a desirable emotion that isn’t necessarily intense, but nevertheless, optimistic. It is often felt when an Upmood user is in a comfortable state with moderate amounts of mood surge.

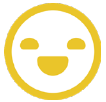

**Happy** is a coveted disposition of lighthearted joy, Upmood will show this emotion when the user has consistently felt an upward spiral of emotional pattern for a certain period of time, such as glee, contentment, and bliss to name a few.

#### Emoji Explanation

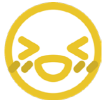

**Excitement** is a transcendent emotion that is indicative of an active behavior. Although, excitement can be borne out of not just positive instances, but also of negative feelings, specifically when an Upmood user is experiencing stress level that is higher than usual.

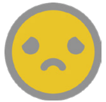

**Unpleasant** emotion appears on Upmood when a user is experiencing mild sense of emotions, mostly negative ones, that can affect the emotional pattern as well as stress levels.

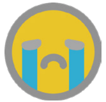

**Sadness** is an overwhelming feeling associated with grief, disappointment, frailty, and misfortune, but Upmood also considers immense pressure, physical tension, and moderate stress levels as triggers of this emotion.

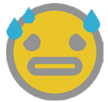

**Anxiety** is an inherent reaction towards stress, making the mind uneasy and nervous of what’s to come. This results to higher stress levels and an arousal on negative emotion of an Upmood user for a period of time.

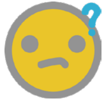

When things don’t make sense or doesn’t translate to what is held to be true, it leads an Upmood user to a state of **confusion**, feeling lost and uncertainty of what to do next. This affects the user’s stress levels and prompts negative emotional arousal.

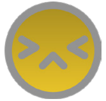

An emotion that is often misconstrued with confused, an Upmood user who has **challenged** for an emotion isn’t necessarily disconcerted but has difficulty to focus, causing the user to feel different kinds of emotion at the same time.
