## Supplementary material for "Between-subject correlation of heart rate variability predicts movie preferences": S2 File

HKU PSY Questionnaire participant no. ___________

Audience Psychology:

**Measuring Emotional Response of Audience based on Heart Rate Variability**

Part A:

Gender: M / F Age: ___________ Occupation: _________________

Phone: iPhone / Android (pls specify_________________________)

Part B:

1. What is your overall rating of the clip (s)
   (1 is the worst while 10 is the best) = _______________________
2. Which plot/part
   a. is the happiest? ___________________________________________
   b. is the saddest? ____________________________________________
   c. is the most memorable? _____________________________________
   d. is the most boring? ________________________________________

3. Will you recommend this compilation of CM to the others? Yes / No
4. Have you watched this clip before? Yes / No

Part C:

1. What is your overall rating of the clip (s)
   (1 is the worst while 10 is the best) = ________________________

2. Which plot/part is
   a. the happiest? ______________________________________________
   b. Is the saddest? ____________________________________________
   c. is the most memorable? _____________________________________
   d. is the most boring? ________________________________________

3. Will you recommend this movie to the others? Yes / No
4. Have you watched this clip before? Yes / No

Part D:

1. What is your overall rating of the clip (s)
   (1 is the worst while 10 is the best) = ________________________

2. Which plot/part is
   a. the happiest? ______________________________________________
   b. the saddest? _______________________________________________
   c. the most memorable? ________________________________________
   d. the most boring? ___________________________________________

3. Will you recommend this movie to the others? Yes / No
4. Have you watched this clip before? Yes / No

Part E:

1. What is your overall rating of the clip (s)
   (1 is the worst while 10 is the best) = ________________________

2. Which plot/part is
   a. the most exciting? __________________________________________
   b. the happiest _______________________________________________
   c. the saddest? _______________________________________________
   d. the most memorable? ________________________________________
   e. the most boring? ___________________________________________

3. Will you recommend this movie to the others? Yes / No
4. Have you watched this clip before? Yes / No

Thank you for your participation, pls return this questionnaire to the researcher
