## Supplementary material for "Between-subject correlation of heart rate variability predicts movie preferences": S3 File

Correlation Matrix

| | | 01 | 02 | 03 | 05 | 08 | 09 | 10 | 11 | 12 | 13 | 14 | 16 | 17 | 18 | 19 | 20 | 21 |
|---|---|---|---|---|---|---|---|---|---|---|---|---|---|---|---|---|---|---|
| 01 | Pearson's r | — | | | | | | | | | | | | | | | | |
| | p-value | — | | | | | | | | | | | | | | | | |
| | N | — | | | | | | | | | | | | | | | | |
| 02 | Pearson's r | 0.013 ** | — | | | | | | | | | | | | | | | |
| | p-value | 0.002 | — | | | | | | | | | | | | | | | |
| | N | 55185 | — | | | | | | | | | | | | | | | |
| 03 | Pearson's r | 0.068 *** | 0.064 *** | — | | | | | | | | | | | | | | |
| | p-value | <.001 | <.001 | — | | | | | | | | | | | | | | |
| | N | 55185 | 55185 | — | | | | | | | | | | | | | | |
| 05 | Pearson's r | 0.058 *** | 0.041 *** | 0.048 *** | — | | | | | | | | | | | | | |
| | p-value | <.001 | <.001 | <.001 | — | | | | | | | | | | | | | |
| | N | 55185 | 55185 | 55185 | — | | | | | | | | | | | | | |
| 08 | Pearson's r | 0.029 *** | 0.045 *** | 0.068 *** | 0.064 *** | — | | | | | | | | | | | | |
| | p-value | <.001 | <.001 | <.001 | <.001 | — | | | | | | | | | | | | |
| | N | 55185 | 55185 | 55185 | 55185 | — | | | | | | | | | | | | |
| 09 | Pearson's r | 0.063 *** | 0.043 *** | 0.047 *** | 0.100 *** | 0.048 *** | — | | | | | | | | | | | |
| | p-value | <.001 | <.001 | <.001 | <.001 | <.001 | — | | | | | | | | | | | |
| | N | 55185 | 55185 | 55185 | 55185 | 55185 | — | | | | | | | | | | | |
| 10 | Pearson's r | 0.054 *** | 0.046 *** | 0.057 *** | 0.062 *** | 0.118 *** | 0.080 *** | — | | | | | | | | | | |
| | p-value | <.001 | <.001 | <.001 | <.001 | <.001 | <.001 | — | | | | | | | | | | |
| | N | 55185 | 55185 | 55185 | 55185 | 55185 | 55185 | — | | | | | | | | | | |
| 11 | Pearson's r | -0.074 | -0.017 | -0.062 | -0.014 | -0.031 | -0.108 | -0.063 | — | | | | | | | | | |
| | p-value | 1.000 | 1.000 | 1.000 | 0.999 | 1.000 | 1.000 | 1.000 | — | | | | | | | | | |
| | N | 55185 | 55185 | 55185 | 55185 | 55185 | 55185 | 55185 | — | | | | | | | | | |
| 12 | Pearson's r | 0.088 *** | 0.027 *** | 0.051 *** | 0.060 *** | 0.082 *** | 0.023 *** | 0.045 *** | -0.029 | — | | | | | | | | |
| | p-value | <.001 | <.001 | <.001 | <.001 | <.001 | <.001 | <.001 | 1.000 | — | | | | | | | | |
| | N | 55185 | 55185 | 55185 | 55185 | 55185 | 55185 | 55185 | 55185 | — | | | | | | | | |
| 13 | Pearson's r | 0.026 *** | 0.024 *** | 0.012 ** | 0.019 *** | -0.001 | 0.006 | 0.006 | 0.026 *** | 0.047 *** | — | | | | | | | |
| | p-value | <.001 | <.001 | 0.003 | <.001 | 0.577 | 0.095 | 0.095 | <.001 | <.001 | — | | | | | | | |
| | N | 55185 | 55185 | 55185 | 55185 | 55185 | 55185 | 55185 | 55185 | 55185 | — | | | | | | | |
| 14 | Pearson's r | 0.019 *** | -0.013 | -0.011 | -0.007 | -0.002 | -0.043 | -0.033 | 0.008 * | 0.010 * | 0.023 *** | — | | | | | | |
| | p-value | <.001 | 0.999 | 0.995 | 0.942 | 0.717 | 1.000 | 1.000 | 0.027 | 0.013 | <.001 | — | | | | | | |
| | N | 55185 | 55185 | 55185 | 55185 | 55185 | 55185 | 55185 | 55185 | 55185 | 55185 | — | | | | | | |
| 16 | Pearson's r | 0.053 *** | -0.004 | 0.012 ** | 0.001 | -0.029 | 0.009 * | -0.002 | -0.007 | -0.009 | 0.001 | -0.005 | — | | | | | |
| | p-value | <.001 | 0.815 | 0.002 | 0.442 | 1.000 | 0.015 | 0.713 | 0.956 | 0.985 | 0.375 | 0.880 | — | | | | | |
| | N | 55185 | 55185 | 55185 | 55185 | 55185 | 55185 | 55185 | 55185 | 55185 | 55185 | 55185 | — | | | | | |
| 17 | Pearson's r | -0.020 | 0.005 | -0.042 | -0.024 | -0.042 | -0.013 | -0.014 | 0.019 *** | -0.020 | 0.027 *** | 0.024 *** | 0.012 ** | — | | | | |
| | p-value | 1.000 | 0.116 | 1.000 | 1.000 | 1.000 | 0.999 | 0.999 | <.001 | 1.000 | <.001 | <.001 | 0.003 | — | | | | |
| | N | 55185 | 55185 | 55185 | 55185 | 55185 | 55185 | 55185 | 55185 | 55185 | 55185 | 55185 | 55185 | — | | | | |
| 18 | Pearson's r | -0.037 | 0.068 *** | 0.042 *** | 0.018 *** | 0.042 *** | -0.022 | 0.020 *** | -0.019 | 0.014 *** | 0.028 *** | 0.017 *** | -0.001 | 0.011 ** | — | | | |
| | p-value | 1.000 | <.001 | <.001 | <.001 | <.001 | 1.000 | <.001 | 1.000 | <.001 | <.001 | <.001 | 0.607 | 0.005 | — | | | |
| | N | 55185 | 55185 | 55185 | 55185 | 55185 | 55185 | 55185 | 55185 | 55185 | 55185 | 55185 | 55185 | 55185 | — | | | |
| 19 | Pearson's r | 0.041 *** | 0.049 *** | 0.057 *** | 0.058 *** | 0.051 *** | 0.064 *** | 0.066 *** | -0.029 | 0.033 *** | 0.053 *** | -0.027 | -0.012 | -0.025 | 0.060 *** | — | | |
| | p-value | <.001 | <.001 | <.001 | <.001 | <.001 | <.001 | <.001 | 1.000 | <.001 | <.001 | 1.000 | 0.997 | 1.000 | <.001 | — | | |
| | N | 55185 | 55185 | 55185 | 55185 | 55185 | 55185 | 55185 | 55185 | 55185 | 55185 | 55185 | 55185 | 55185 | 55185 | — | | |
| 20 | Pearson's r | 0.051 *** | 0.054 *** | 0.056 *** | 0.070 *** | 0.001 | 0.054 *** | 0.040 *** | -0.037 | 0.038 *** | 0.018 *** | 0.013 *** | -0.004 | -0.001 | 0.029 *** | 0.046 *** | — | |
| | p-value | <.001 | <.001 | <.001 | <.001 | 0.377 | <.001 | <.001 | 1.000 | <.001 | <.001 | <.001 | 0.819 | 0.633 | <.001 | <.001 | — | |
| | N | 55185 | 55185 | 55185 | 55185 | 55185 | 55185 | 55185 | 55185 | 55185 | 55185 | 55185 | 55185 | 55185 | 55185 | 55185 | — | |
| 21 | Pearson's r | 0.079 *** | 0.013 ** | 0.004 | 0.036 *** | 0.011 ** | 0.032 *** | 0.011 ** | -0.058 | 0.036 *** | -0.055 | 0.005 | 0.013 ** | 0.019 *** | -0.015 | -0.028 | 0.029 *** | — |
| | p-value | <.001 | 0.001 | 0.193 | <.001 | 0.005 | <.001 | 0.006 | 1.000 | <.001 | 1.000 | 0.134 | 0.001 | <.001 | 1.000 | 1.000 | <.001 | — |
| | N | 55185 | 55185 | 55185 | 55185 | 55185 | 55185 | 55185 | 55185 | 55185 | 55185 | 55185 | 55185 | 55185 | 55185 | 55185 | 55185 | — |

*Note.* $H_a$ is positive correlation

*Note.* * p < .05, ** p < .01, *** p < .001, one-tailed
