## Supplementary material for "Between-subject correlation of heart rate variability predicts movie preferences": S4 File

### DASS21

Name: Date:

Please read each statement and circle a number 0, 1, 2 or 3 which indicates how much the statement applied to you **over the past week**. There are no right or wrong answers. Do not spend too much time on any statement.

The rating scale is as follows:

0 Did not apply to me at all
1 Applied to me to some degree, or some of the time
2 Applied to me to a considerable degree or a good part of time
3 Applied to me very much or most of the time

| | | | | | |
|---|---|---|---|---|---|
| 1 (s) | I found it hard to wind down | 0 | 1 | 2 | 3 |
| 2 (a) | I was aware of dryness of my mouth | 0 | 1 | 2 | 3 |
| 3 (d) | I couldn’t seem to experience any positive feeling at all | 0 | 1 | 2 | 3 |
| 4 (a) | I experienced breathing difficulty (e.g. excessively rapid breathing, breathlessness in the absence of physical exertion) | 0 | 1 | 2 | 3 |
| 5 (d) | I found it difficult to work up the initiative to do things | 0 | 1 | 2 | 3 |
| 6 (s) | I tended to over-react to situations | 0 | 1 | 2 | 3 |
| 7 (a) | I experienced trembling (e.g. in the hands) | 0 | 1 | 2 | 3 |
| 8 (s) | I felt that I was using a lot of nervous energy | 0 | 1 | 2 | 3 |
| 9 (a) | I was worried about situations in which I might panic and make a fool of myself | 0 | 1 | 2 | 3 |
| 10 (d) | I felt that I had nothing to look forward to | 0 | 1 | 2 | 3 |
| 11 (s) | I found myself getting agitated | 0 | 1 | 2 | 3 |
| 12 (s) | I found it difficult to relax | 0 | 1 | 2 | 3 |
| 13 (d) | I felt down-hearted and blue | 0 | 1 | 2 | 3 |
| 14 (s) | I was intolerant of anything that kept me from getting on with what I was doing | 0 | 1 | 2 | 3 |
| 15 (a) | I felt I was close to panic | 0 | 1 | 2 | 3 |
| 16 (d) | I was unable to become enthusiastic about anything | 0 | 1 | 2 | 3 |
| 17 (d) | I felt I wasn’t worth much as a person | 0 | 1 | 2 | 3 |
| 18 (s) | I felt that I was rather touchy | 0 | 1 | 2 | 3 |
| 19 (a) | I was aware of the action of my heart in the absence of physical exertion (e.g. sense of heart rate increase, heart missing a beat) | 0 | 1 | 2 | 3 |
| 20 (a) | I felt scared without any good reason | 0 | 1 | 2 | 3 |
| 21 (d) | I felt that life was meaningless | 0 | 1 | 2 | 3 |

### DASS-21 Scoring Instructions

The DASS-21 should not be used to replace a face to face clinical interview. If you are experiencing significant emotional difficulties you should contact your GP for a referral to a qualified professional.

**Depression, Anxiety and Stress Scale - 21 Items (DASS-21)**

The Depression, Anxiety and Stress Scale - 21 Items (DASS-21) is a set of three self-report scales designed to measure the emotional states of depression, anxiety and stress.

Each of the three DASS-21 scales contains 7 items, divided into subscales with similar content. The depression scale assesses dysphoria, hopelessness, devaluation of life, self-deprecation, lack of interest / involvement, anhedonia and inertia. The anxiety scale assesses autonomic arousal, skeletal muscle effects, situational anxiety, and subjective experience of anxious affect. The stress scale is sensitive to levels of chronic non-specific arousal. It assesses difficulty relaxing, nervous arousal, and being easily upset / agitated, irritable / over-reactive and impatient. Scores for depression, anxiety and stress are calculated by summing the scores for the relevant items.

The DASS-21 is based on a dimensional rather than a categorical conception of psychological disorder. The assumption on which the DASS-21 development was based (and which was confirmed by the research data) is that the differences between the depression, anxiety and the stress experienced by normal subjects and clinical populations are essentially differences of degree. The DASS-21 therefore has no direct implications for the allocation of patients to discrete diagnostic categories postulated in classificatory systems such as the DSM and ICD.

Recommended cut-off scores for conventional severity labels (normal, moderate, severe) are as follows:

NB Scores on the DASS-21 will need to be multiplied by 2 to calculate the final score.

| | **Depression** | **Anxiety** | **Stress** |
|---|---|---|---|
| Normal | 0-9 | 0-7 | 0-14 |
| Mild | 10-13 | 8-9 | 15-18 |
| Moderate | 14-20 | 10-14 | 19-25 |
| Severe | 21-27 | 15-19 | 26-33 |
| Extremely Severe | 28+ | 20+ | 34+ |

Lovibond, S.H. & Lovibond, P.F. (1995). Manual for the Depression Anxiety & Stress Scales. (2nd Ed.)Sydney: Psychology Foundation.
