## Supplementary material for "Between-subject correlation of heart rate variability predicts movie preferences": S5 File

### The Quick Inventory of Depressive Symptomatology (16-Item) (Self-Report) (QIDS-SR16)

Name or ID: ______________________________ Date: ______________________________

**CHECK THE ONE RESPONSE TO EACH ITEM THAT BEST DESCRIBES YOU FOR THE PAST SEVEN DAYS.**

#### During the past seven days...

**1. Falling Asleep:**

☐ 0 I never take longer than 30 minutes to fall asleep.

☐ 1 I take at least 30 minutes to fall asleep, less than half the time.

☐ 2 I take at least 30 minutes to fall asleep, more than half the time.

☐ 3 I take more than 60 minutes to fall asleep, more than half the time.

**2. Sleep During the Night**

☐ 0 I do not wake up at night.

☐ 1 I have a restless, light sleep with a few brief awakenings each night.

☐ 2 I wake up at least once a night, but I go back to sleep easily.

☐ 3 I awaken more than once a night and stay awake for 20 minutes or more, more than half the time.

**3. Waking Up Too Early:**

☐ 0 Most of the time, I awaken no more than 30 minutes before I need to get up.

☐ 1 More than half the time, I awaken more than 30 minutes before I need to get up.

☐ 2 I almost always awaken at least one hour or so before I need to, but I go back to sleep eventually.

☐ 3 I awaken at least one hour before I need to, and can't go back to sleep.

**4. Sleeping Too Much:**

☐ 0 I sleep no longer than 7-8 hours/night, without napping during the day.

☐ 1 I sleep no longer than 10 hours in a 24-hour period including naps.

☐ 2 I sleep no longer than 12 hours in a 24-hour period including naps.

☐ 3 I sleep longer than 12 hours in a 24-hour period including naps.

#### During the past seven days...

**5. Feeling Sad:**

☐ 0 I do not feel sad.

☐ 1 I feel sad less than half the time.

☐ 2 I feel sad more than half the time.

☐ 3 I feel sad nearly all of the time.

#### Please complete either 6 or 7 (not both)

**6. Decreased Appetite:**

☐ 0 There is no change in my usual appetite.

☐ 1 I eat somewhat less often or lesser amounts of food than usual.

☐ 2 I eat much less than usual and only with personal effort.

☐ 3 I rarely eat within a 24-hour period, and only with extreme personal effort or when others persuade me to eat.

**- OR -**

**7. Increased Appetite:**

☐ 0 There is no change from my usual appetite.

☐ 1 I feel a need to eat more frequently than usual.

☐ 2 I regularly eat more often and/or greater amounts of food than usual.

☐ 3 I feel driven to overeat both at mealtime and between meals.

#### Please complete either 8 or 9 (not both)

**8. Decreased Weight (Within the Last Two Weeks):**

☐ 0 I have not had a change in my weight.

☐ 1 I feel as if I have had a slight weight loss.

☐ 2 I have lost 2 pounds or more.

☐ 3 I have lost 5 pounds or more.

**- OR -**

**9. Increased Weight (Within the Last Two Weeks):**

☐ 0 I have not had a change in my weight.

☐ 1 I feel as if I have had a slight weight gain.

☐ 2 I have gained 2 pounds or more.

☐ 3 I have gained 5 pounds or more.

#### The Quick Inventory of Depressive Symptomatology (16-Item) (Self-Report) (QIDS-SR16)

##### During the past seven days...

**10. Concentration / Decision Making:**

- ☐ 0 There is no change in my usual capacity to concentrate or make decisions.
- ☐ 1 I occasionally feel indecisive or find that my attention wanders.
- ☐ 2 Most of the time, I struggle to focus my attention or to make decisions.
- ☐ 3 I cannot concentrate well enough to read or cannot make even minor decisions.

**11. View of Myself:**

- ☐ 0 I see myself as equally worthwhile and deserving as other people.
- ☐ 1 I am more self-blaming than usual.
- ☐ 2 I largely believe that I cause problems for others.
- ☐ 3 I think almost constantly about major and minor defects in myself.

**12. Thoughts of Death or Suicide:**

- ☐ 0 I do not think of suicide or death.
- ☐ 1 I feel that life is empty or wonder if it's worth living.
- ☐ 2 I think of suicide or death several times a week for several minutes.
- ☐ 3 I think of suicide or death several times a day in some detail, or I have made specific plans for suicide or have actually tried to take my life.

**13. General Interest**

- ☐ 0 There is no change from usual in how interested I am in other people or activities.
- ☐ 1 I notice that I am less interested in people or activities.
- ☐ 2 I find I have interest in only one or two of my formerly pursued activities.
- ☐ 3 I have virtually no interest in formerly pursued activities.

##### During the past seven days...

**14. Energy Level:**

- ☐ 0 There is no change in my usual level of energy.
- ☐ 1 I get tired more easily than usual.
- ☐ 2 I have to make a big effort to start or finish my usual daily activities (for example, shopping, homework, cooking, or going to work).
- ☐ 3 I really cannot carry out most of my usual daily activities because I just don't have the energy.

**15. Feeling Slowed Down:**

- ☐ 0 I think, speak, and move at my usual rate of speed.
- ☐ 1 I find that my thinking is slowed down or my voice sounds dull or flat.
- ☐ 2 It takes me several seconds to respond to most questions and I'm sure my thinking is slowed.
- ☐ 3 I am often unable to respond to questions without extreme effort.

**16. Feeling Restless:**

- ☐ 0 I do not feel restless.
- ☐ 1 I'm often fidgety, wringing my hands, or need to shift how I am sitting.
- ☐ 2 I have impulses to move about and am quite restless.
- ☐ 3 At times, I am unable to stay seated and need to pace around.
